# Changes in the Public IgM Repertoire and Its Idiotypic Connectivity in Alzheimer’s Disease and Frontotemporal Dementia

**DOI:** 10.1101/2024.07.15.603559

**Authors:** Shina Pashova-Dimova, Peter Petrov, Sena Karachanak-Yankova, Diana Belezhanska, Yavor Zhelev, Shima Mehrabian, Draga Toncheva, Lachezar Traykov, Anastas Pashov

## Abstract

Alzheimer’s disease (AD) and frontotemporal dementia (FTD) are prevalent neurodegenerative disorders. Early diagnosis is challenging due to the lack of definitive biomarkers and reliance on invasive procedures. Immune biomarkers, particularly those reflecting the interaction between the central nervous system (CNS) and the peripheral immune system, have shown promise for non-invasive detection through blood samples. This study investigates the reactivity of serum IgM and IgG from AD and FTD patients against a library of mimotopes representing public IgM reactivities in healthy donors. Serum samples from AD, FTD, and other neurodegenerative dementias (ND) and controls were tested on peptide microarrays. The samples were pooled to mitigate individual variability. The reactivity data were analyzed using graphs to represent the cross-reactivity networks. The analysis revealed distinct reactivity patterns for the studied groups. Public IgM reactivities showed significant correlations with neurodegenerative conditions, with AD and FTD exhibiting loss or gain of specific IgM reactivities. Graph analysis highlighted significant differences between disease and control groups in graph density, clustering, and assortativity parameters. Mimotopes of IgM reactivities lost in dementia, particularly in AD, exhibited significant homology to HCDR3 sequences of human antibodies. Furthermore, clusters of reactivities showed significant distinctions between AD and FTD, with IgG reactivities providing additional differentiation. Several self-proteins related to neurodegeneration proved to have sequences homologous to disease-associated mimotopes. Interestingly, the beta-propeller signature sequence YWTD found in ApoE’s receptor LRP1 proved a characteristic epitope for IgG in FTD but not AD. At the same time, the respective public |gM mimotope YWTDSSR coincides with a highly conserved sequence in many microorganisms and sequences found in human HCDR3. Thus, the public IgM repertoire, characterized by its broad reactivity and inherent autoreactivity, offers valuable insights into the immunological alterations in neurodegenerative diseases. The study supports the potential of IgM and IgG reactivity profiles as another compartment of non-invasive biomarkers for early diagnosis and differentiating AD and FTD.

## 1. Introduction

Alzheimer’s disease (AD) and frontotemporal dementia (FTD) are among the most prevalent neurodegenerative disorders affecting the elderly population (Musa et al., 2020). Even the basic pathogenic mechanisms of these diseases are still debated (Castellani et al., 2022) But it is clear that they involve complex system-level interactions, including immunological mechanisms (Chen and Colonna, 2022). A better understanding of the pathogenesis of neurodegenerative disease serves the development of potential therapy approaches and better and earlier diagnosis. The diagnosis of AD and FTD remains a significant challenge due to the lack of definitive biomarkers and the need for invasive procedures like cerebrospinal fluid (CSF) analysis or positron emission tomography (PET) scans (Gifford et al., 2023; Shue et al., 2024). Recent advancements in plasma biomarkers and computational approaches have opened up new avenues for the early diagnosis of these neurodegenerative diseases. Immune biomarkers, particularly those related to the cross-talk between the central nervous system (CNS) and the immune system, have emerged as promising tools for the detection of AD and FTD (Krix et al., 2024). These serum biomarkers are more accessible and cost-effective than traditional methods, and their diversity justifies the use of machine learning approaches, combining them with further information (e.g., age, sex, APOE genotype) into predictive models supporting an earlier diagnosis.

Among the promising immunological biomarkers, antibodies have a unique role because of their highly diverse repertoire of specificities. Previous studies have demonstrated that patients with AD exhibit unique IgG antibody binding patterns compared to healthy controls and those with other types of dementia. For instance, Restrepo et al. (2011) explored the application of a peptide microarray containing 10,000 random sequence peptides to profile the antibody repertoire of AD patients and transgenic mouse models. The study found that AD patients and mice with cerebral amyloidosis displayed distinct immunoprofiles, suggesting the utility of the immunosignature technique in AD diagnosis (Restrepo et al., 2011). Some sets of IgG autoantibodies with better-defined autoantigen specificities have shown great promise for the development of multi-disease biomarkers, including for the early detection of AD (DeMarshall et al., 2017; DeMarshall et al., 2023; Nagele et al., 2011). Unlike the highly specific IgG, which is usually the product of an adaptive immune response, the IgM repertoire abounds in constitutively produced natural antibody, and at least 20% of them are physiologically autoreactive (Imkeller and Wardemann, 2018). Since it is also more diverse than its IgG counterpart and better overlapping with the public reactivities (Galson et al., 2015). Thus, the individual IgM repertoire is better poised as a sensor of the internal environment before the development of overt autoimmunity. With this working hypothesis, we started developing a diagnostic platform applicable to a very broad range of pathologies that is based on applying IgOme approaches (Ryvkin et al., 2012) for extracting patterns of diagnostic public IgM reactivities (Ferdinandov et al., 2023; Pashov et al., 2019; Pashova-Dimova et al., 2023; Pashova et al., 2022; Shivarov et al., 2020).

In this manuscript, we present our findings on the reactivity of serum IgM and IgG from AD and FTD patients to a large library of mimotopes of the public IgM reactivities found in healthy donors. In addition, the cross-reactivities to the same mimotopes in the IgG repertoire were tested, hoping to estimate the degree of the adaptive immune response contribution to the observed binding patterns. We focus on the differences in antibody reactivity graphs generated from the binding data as a formalism for cross-reactivity networks. Somewhat surprisingly, losing or gaining public IgM reactivities in AD and FTD correlated with their cross-reactivity with human immunoglobulin HCDR3 sequences seen as potential idiotopes.

## 2. Materials and methods

### 2.1. Patients and sera

Patients were recruited at the Clinic of Neurology, Medical University - Sofia. The diagnosis of AD was made according to the NIA-AA criteria (McKhann et al., 2011). A diagnosis of behavioral variant of FTD and primary progressive aphasia (PPA) was considered based on the current revised diagnostic guidelines (Gorno-Tempini et al., 2011; Rascovsky et al., 2011). A group of patients with other neurodegenerative diseases (Lewy body disease, Parkinson’s disease with dementia, and Huntington disease) was also included. A summary of the patients’ clinical data is presented in Table 1. Sera were collected from three groups of 8 patients diagnosed with AD, FTD, or other neurodegenerative dementias and compared to sera from 6 sex and age-matched control group patients without dementia (C). All participants or their caregivers gave written informed consent to participation in the study. The informed consent forms and study protocols were approved by the local ethics committee, UH “Alexandrovska”, and performed in harmony with the Helsinki Declaration.

**Table 1.**
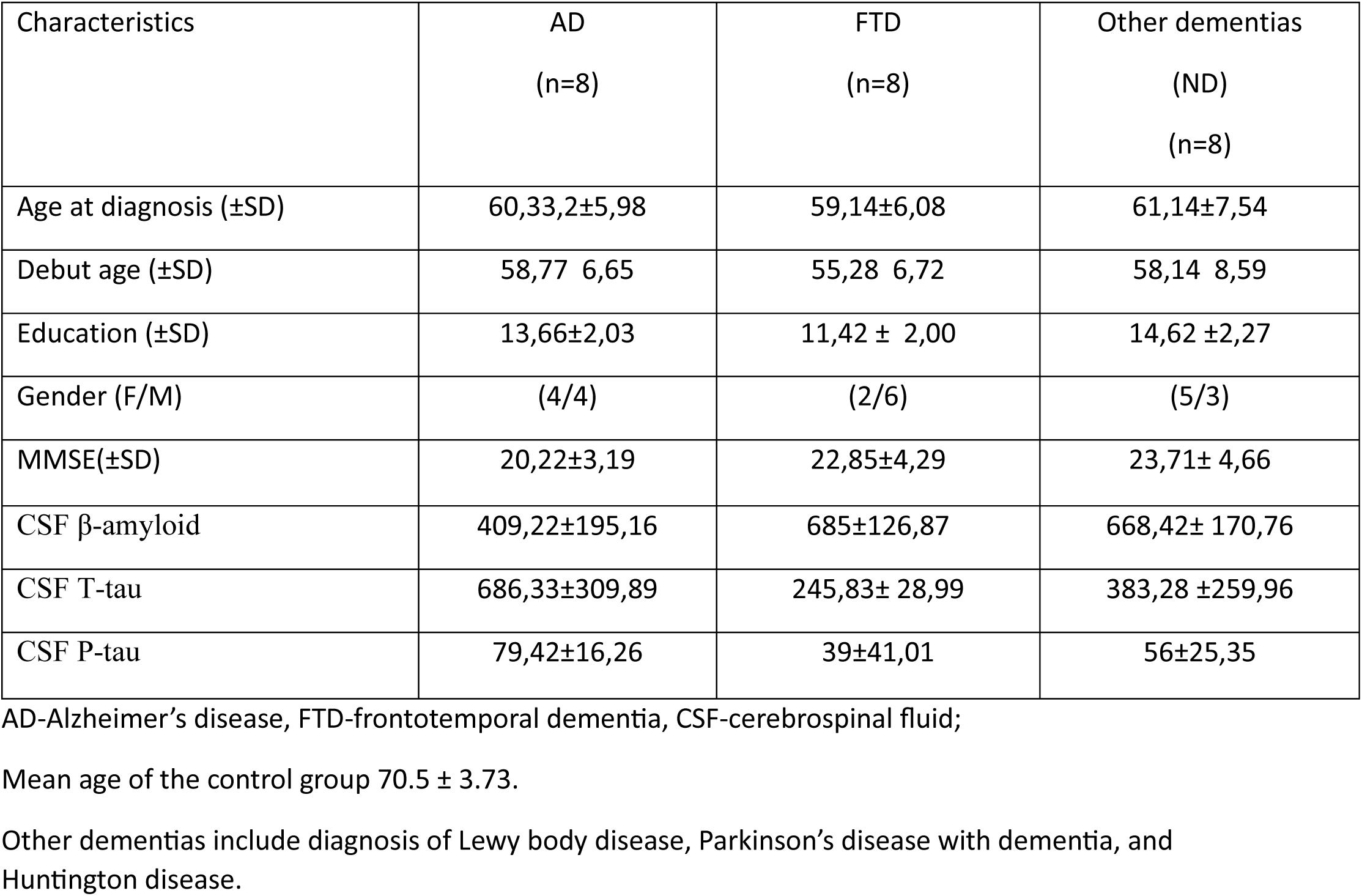
Summary of the clinical data of the patients included.

### 2.2. IgM and IgG isolation

The serum samples were thawed and incubated at 37°C for 30 min to dissolve the IgM aggregates. After spinning for 10 min at 20,000 × g, the IgM concentration was determined by ELISA, and the sera were diluted to 10 µg/ml IgM. The samples used in the study were limited to 5 per diagnosis, so some of the samples had to be pooled. Pooling creates correlation between the samples, which makes cross-validation and machine learning impossible. Since this limitation was inevitable, it was decided that all samples would be pools of two patients’ appropriately diluted sera as an intermediate averaging of reactivities in the groups and giving a slight advantage to public reactivities. The pooling schemes in terms of serum number were: AD – 1+6, 2+7, 3+8, 4+8, and 5+2, FTD - 1+6, 2+7, 3+8, 4+1, and 5+2, ND - 1+6, 2+7, 3+8, 4+1, and 5+2, and C – 1+3, 2+6, 3+4, 4+2, 5+1.

The patients’ blood groups were detected by reverse blood group typing of the sera using the test erythrocytes kit Biotestcell (BioRad).

### 2.3. Peptide microarray binding assay

#### 2.3.1. Sequence selection

Previously, we have studied a library of 2×10^5^ peptide mimotopes of the public IgM of healthy donors (Pashov et al., 2019). The selection of public reactivities is assured by using IgM isolated from a blood plasma pool from 10 000 individuals, which substantially dilutes the private IgM specificities. Using a Gibbs sampling approach (Andreatta et al., 2013) (https://services.healthtech.dtu.dk/services/GibbsCluster-2.0/), the sequences were clustered optimally in 790 clusters (Pashov et al., 2019). Here, we further analyzed this grouping by constructing a graph of the clusters. For each primary cluster, the neighboring clusters were determined based on sequence proximity. The latter was based on the next best cluster information for each of the sequences of a given cluster as reported by the algorithm. If a cluster had more than n sequences close to the same alternative cluster, the two clusters were connected by an edge. Most (675) of the clusters remained disconnected, but some formed the graph components connecting 2 to 10 clusters. The threshold n was adjusted to produce fewer than 730 components (a number defined by the peptide array format). These components were considered the secondary clusters (n=726). The singletons in this graph represent primary clusters, and the Gibbs Sampling algorithm provided the most relevant representative sequence from each cluster. Each of the secondary clusters was converted to an IgOme graph of sequences (Pashova et al., 2022), and the vertex with the highest eigencentrality was used to define the representative sequence. In this way, a set of 726 peptides that sampled symmetrically the space of the 7-mer mimotopes of the public IgM reactivities was selected and used to synthesize the peptide arrays.

#### 2.3.2. Peptide microarrays design

PEPperPRINT^TM^ (Heidelberg, Germany) produced custom peptide microarrays with 726 7 amino acid residues long linear peptides (see 2.3.1). The peptides were synthesized in situ and attached to the functionalized chip surface by their C-terminus and through a GSGSG spacer. The layout of the microarrays contained the respective peptide spots duplicated at random positions.

Supplementary Table 1 shows the sequences of the peptides used. The peptide arrays were blocked for 60 min using PBS, pH 7.4, and 0.05% Tween 20 with 1% BSA with constant shaking; washed 3 × 1 min with PBS, pH 7.4, and 0.05% Tween 20 followed by incubation with sera in dilutions equivalent to 0.01 mg/mL IgM (established by determining the immunoglobulin concentrations by ELISA) on an orbital shaker overnight at 4°C. After washing 3 × 1 min, the arrays were incubated with secondary antibodies at room temperature, washed, rinsed with distilled water, and dried by spinning vertically in empty 50 mL test tubes at 100×g for 2 min. Two different secondary antibodies were used to measure simultaneously the binding of IgG and IgM at wavelengths of detection, respectively, 544 nm and 647 nm.

### 2.4. Microarray data analysis

#### 2.4.1. Acquisition, cleaning and normalization

The Innoscan 1100 (Innopsys, Carbonne, France) microarray scanner was used to acquire the stained microarray fluorescence images. MAPIX software was used for the subsequent densitometry. The analysis of the densitometry data was performed using publicly available packages from the R statistical environment (*Bioconductor, Biostrings, limma, pepStat, igraph, clvalid, gplots, future.apply, FactoMIneR, factoextra, stringi, stringdist, rBLAST, rgl,* etc.) as well as in-house developed R functions. The data and the scripts of the analysis are deposited at https://github.com/ansts/ADFTDIgOme. The cross-reactivity graph strategy was previously described (Ferdinandov et al., 2023). Here it is used with some modifications. Briefly, the raw data was cleaned based on the flags set during densitometry. The background was defined based on the gradients of staining between the randomly positioned duplicate peptide spots and subtracted (Pashov et al., 2019).

The logarithms of the data were normalized relative to the amino acid residue composition using the pepstat Bioconductor package as described previously (Pashov et al., 2019). This compensates for binding, which is dependent on the overall charge and hydrophobicity rather than on the sequence. Further, the data was normalized between arrays using the *limma*::normalizeCyclicLoess function with the “affy” method applied in 3 iterations separately for the IgM and IgG datasets. At the end of this stage, the datasets were based on 722 mimotopes.

#### 2.4.2. Graph representation

Each of the isotype sets was used to generate a separate cross-reactivity graph. The criterion for cross-reactivity between two peptides used to define the edges of the graphs was a similarity measure. It was based on the reciprocal of the Manhattan distance between the respective profiles of reactivities with the 20 samples. Each pair of profiles was first scaled and centered using the median and the interquartile range. If *M* is the n×k matrix of the n reactivity values with k samples and *i*1 and *i*2 are two rows corresponding to two peptides (profiles), the quartiles of each row *M*_*i*_ being

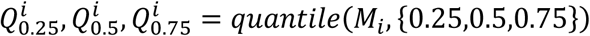

the similarity measure is calculated as:

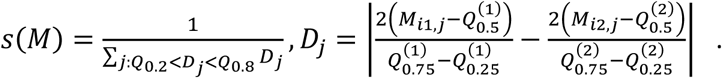

Only the *D* values between the quantiles 0.2 and 0.8 were used. The filtering out of the extreme values of *D* is done to prevent the decision about cross-reactivity from being taken on the basis of just a couple of similar values. The concept of defining cross-reactivity by testing multiple individual repertoires depends on the multiplicity of samples with correlated reactivities. The criterion was tested using the data form (Ferdinandov et al., 2023). The ROC curve and the selected threshold for connecting the vertices of the graphs were analyzed as previously described (Ferdinandov et al., 2023). The threshold selected ensured specificity of 0.9 and sensitivity of 0.6, thus emphasizing only the most probable cross-reactivities (Fig. 1).

**Figure 1.**
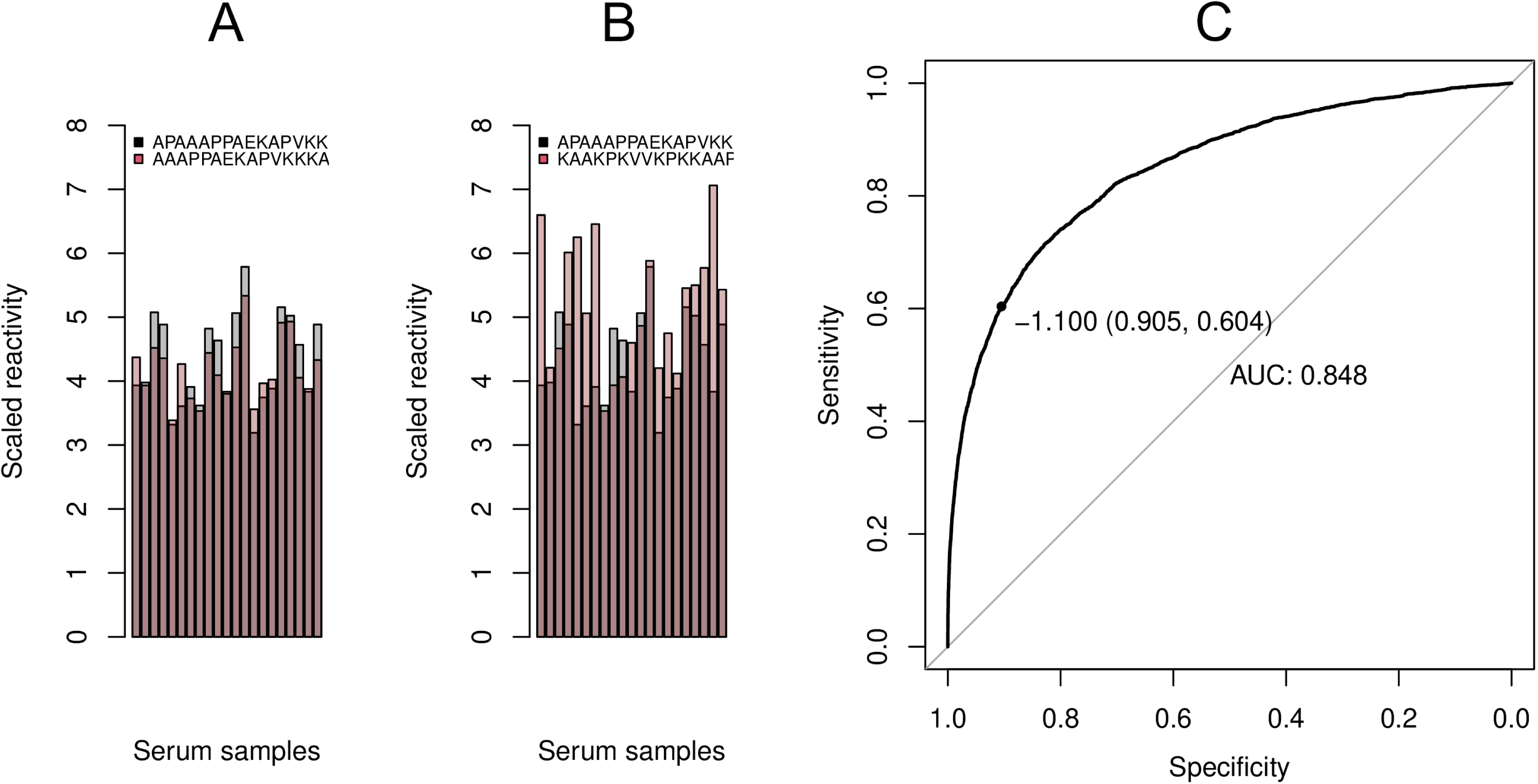
Tuning the similarity measure used to construct the graph. A and B - Examples of profile pairs of cross-reactive (A) and not cross-reactive peptides (B), C - ROC curve illustrating the capacity of the similarity criterion to classify 4150 pairs of peptides overlapping in 11/15 positions as a model of cross-reactive peptide pairs compared to a set of 10,000 pairs of dissimilar peptides sharing a longest common subsequence of fewer than 3/15 position (data from (Ferdinandov et al., 2023)). The tradeoff between sensitivity and specificity chosen log10(s)=-1.1 (specificity = 0.905, sensitivity = 0.604, AUC=0.848).

An edge in a reactivity graph *G* can be interpreted as a cross-reactivity relation between two peptides. Since the cross-reactivity is inferred from similarity of the reactivity profiles across samples, the same peptide can be cross-reactive with other peptides based on different parts of the profile. The respective edges will represent potentially different antibody reactivities. Therefore, the topology of the reactivity graph does not represent the map of cross reactivities with the necessary detail unless the qualitative differences between the edges are figured in. To improve the clustering so that it can be used to formalize the concept of latent antibody reactivities, a second graph – *GE*, was constructed with vertices representing the edges in *G*. *GE* was not a line graph of *G*. The vertices in *GE* (originally edges in *G*) are connected not when they have a common incident vertex (like in a line graph) but when they stand for correlating profiles *D*_j_. Given the reactivity graph *G*, and the raw data matrix *M*, from which *G* is constructed,

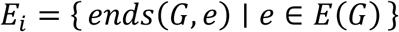

are the endpoints of each edge where *ends*(*G*, *e*) represents the pair of vertices at the ends of the edge *e* in graph *G*. The distance matrix

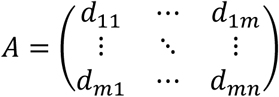

where the distances *d*_{*jk*}_ = |*μ*(*M*[*l*_j_, :]) − *μ*(*M*[*l*_*k*_, :])| and *μ* is the vector of column-wise means of each row pair, *l_j_*and *l_k_* being the two pairs of indices of the respective pairs of profiles *j* and *k*. After setting the diagonal of *A* to zero, this became effectively the adjacency matrix of the *GE* graph. The weights of the edges in this graph (as distances) are given by the elements of *A*. Next, a cluster size optimization was performed by varying the threshold w_*i*_ for edge deletion. The threshold *c*_*t*_ was determined by maximizing the product of the cluster sizes *s_k_* of the clusters larger than 5 vertices:

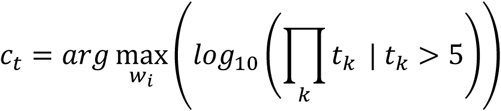

The edges with distances greater than the optimal threshold *c*_*t*_ were deleted. The ultimate form of the graph *GE*, thus obtained, was clustered to obtain clusters of what are respectively edges in the context of the original graph *G*. They were labeled edge clusters. Every edge cluster was defined in terms of the vertices - its end points in the original graph *G*. The final grouping of reactivities was obtained as intersections of the *G* clusters with the vertex clustering derived from the *GE* edge clusters. The groups of reactivity obtained are not a partition (like the clusters) but a cover (they overlap) because different edges can have overlapping vertices as endpoints.

To test the significance of various graph parameters (like size of large component, number of clusters, assortativity of vertex parameters, degree and maximal clique distributions, etc.) the baseline values for these parameters were estimated using 2 sets of 100 random graphs generated using the same algorithm but starting with the scrambled raw data matrices for IgM and IgG reactivities respectively. All graph analysis was performed using the igraph package.

Let 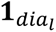 be the indicator function of the diagnosis group dia_*l*_ ∈ {“A”,“C”, “F”, “ND“} and let *r*_*i*_ = [*r*_*i*1_, *r*_*i*1_, …, *r*_*ik*_] represent the *i-th* row (mimotope) of the reactivity data matrix *M*. The rows are ranked: 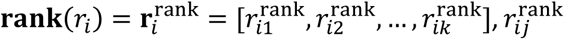 denotes the *j-th* element of the *i-th* row. Then, the sum of ranks for the group *dia*_*l*_ for the *i-th* row is:

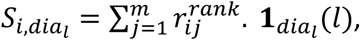

Where 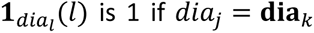 is 1 if *dia*_*j*_ = **dia**_*k*_ and 0 otherwise. The 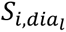 was used to quantify the degree of association of each reactivity with each diagnosis.

For the visualization, the graphs were embedded using the eigenvectors of the graph Laplacian of each isotype graph (spectral embedding). The eigenvectors that corresponded to the k smallest eigenvalues were used. The cutoff to determine k was based on the irregular increase in the ordered eigenvalues (Suppl. Fig. 2). The exact position of the vertices in the two-dimensional projections is derived by UMAP embedding from k dimensions to two dimensions. UMAP was considered appropriate for the final 2D embedding because applying it to a relatively low-dimensional dataset reduces the distortion of distances. The 3D visualization was done with the help of the rgl package. The graph clustering was done using the Leiden algorithm with modularity optimization and resolution parameter 1.

### 2.5. Machine learning model

To mitigate the inter-sample correlation, a specific cross-validation scheme was applied to classify the three diagnoses using a support vector machine-based (SVM) model. The feature selection step included a filtering step removing features with a Mahalanobis distance less than the 0.35 quantile of its distribution in the space of the samples. This was followed by a recursive feature elimination on the reduced feature set (typically about 400 features) using a compound clustering criterion based on Dunn’s, the connectivity, and the Baker-Hubert Gamma criteria as previously described (Pashov et al., 2019). For the cross-validation, to disconnect the correlated pools, sets of samples were left out (1-3 per step), so that they contained no serum sample used in the rest of the pools. Because of the small number of samples, the labeling was only dichotomous and applied separately for AD/FTD vs C/ND, AD/C vs FTD/ND, and FTD/C vs AD/ND. The aim was to assess roughly the observed patterns’ separability and generalizability. The optimized SVM parameters were: gamma = 0.001, and the constant was set at 35. Before training the model, the data was projected on the first six principal components following the n/3 rule for optimal dimensionality reductionrelative to size of the sample, prompted by the peaking phenomenon (Sima and Dougherty, 2008).

### 2.6. External data sources

The human immunoglobulin heavy chain junction regions found in the NCBI protein database (1 012 520 different entries) was used. The junction regions include the D gene product and correspond to the most diverse portion of the HCDR3 (heavy chain complementarity-determining region 3). From them, 4 433 252 unique overlapping 7-residues sequence frames were derived to compare the mimotopes and find potential linear idiotopes among the latter. The comparison to a given mimotope sequence was done by finding the number of HCDR3 region 7-mer sequences within a longest common subsequence (LCS) distance of 2, i.e. – with no more than 1 mismatch. A set of 217 442 7-mer sequences from a previously published analysis (Matochko et al., 2014; Matochko and Derda, 2013) amplified from the same PDL (Ph.D −7, New England Biolabs) without prior selection was used as a background non-selected library to bootstrap the frequency of idiotopes among mimotopes. To perform protein BLAST (using the rBLAST package), the human RefSeq protein database from NCBI was used (136 194 sequences). The filter was set to 1 mismatch. The criterion of editing distance of 2 (one deletion, one insertion) was previously established (Pashova et al., 2022) based on data from the Biopanning data bank (http://i.uestc.edu.cn/bdb/) (He et al., 2016), summarizing results from phage display screening with monoclonal antibodies.

### 2.7. Statistics

In this study, the groups were compared using non-parametric tests – the Wilcoxon rank sum test and Dunn’s test for multiple comparisons with Benjamini-Hochberg (BH) adjustment of the p values where necessary. Principal component analysis (PCA) was used for data exploration and dimensionality reduction using the *FactoMineR* and *factoextra* packages. Each reactivity was compared to the blood group antigen score. The blood group antigen score equaled the number of pooled sera coming from patients expressing that antigen. Thus, it took only 3 values: 0, 1, and 2.

## 3. Results

An array of 722 representative mimotopes of public IgM reactivities in healthy donors (Pashov et al., 2019) were used to probe the repertoires of serum IgM and IgG from patients with neurodegenerative diseases. To this end, samples from AD (n=8), FTD (n=8), ND (n=8), and controls without clinical dementia (n=6) were pooled in pairs to yield four groups of five serum samples. These were tested on the IgM mimotopes in oriented planar peptide microarrays (Suppl. Table 1).

### 3.1. The reactivity data distinguishes diagnoses

After cleaning and normalization, the data from serum IgM and IgG binding to the array of public IgM mimotopes showed considerable correlation between all samples (Suppl. Fig. 1). Amino acid residue composition bias is most probably not the cause because the data was normalized with respect to that. Some part of this correlation may be due to testing public reactivities that reflect constant characteristics of the repertoire. Another contribution may be from the between-array normalization. In the overview of the data analyzed by PCA, the latent variable underlying this correlation was isolated in the first component. Excluding this factor, the other components separated the samples by diagnosis (Fig.2).

**Figure 2.**
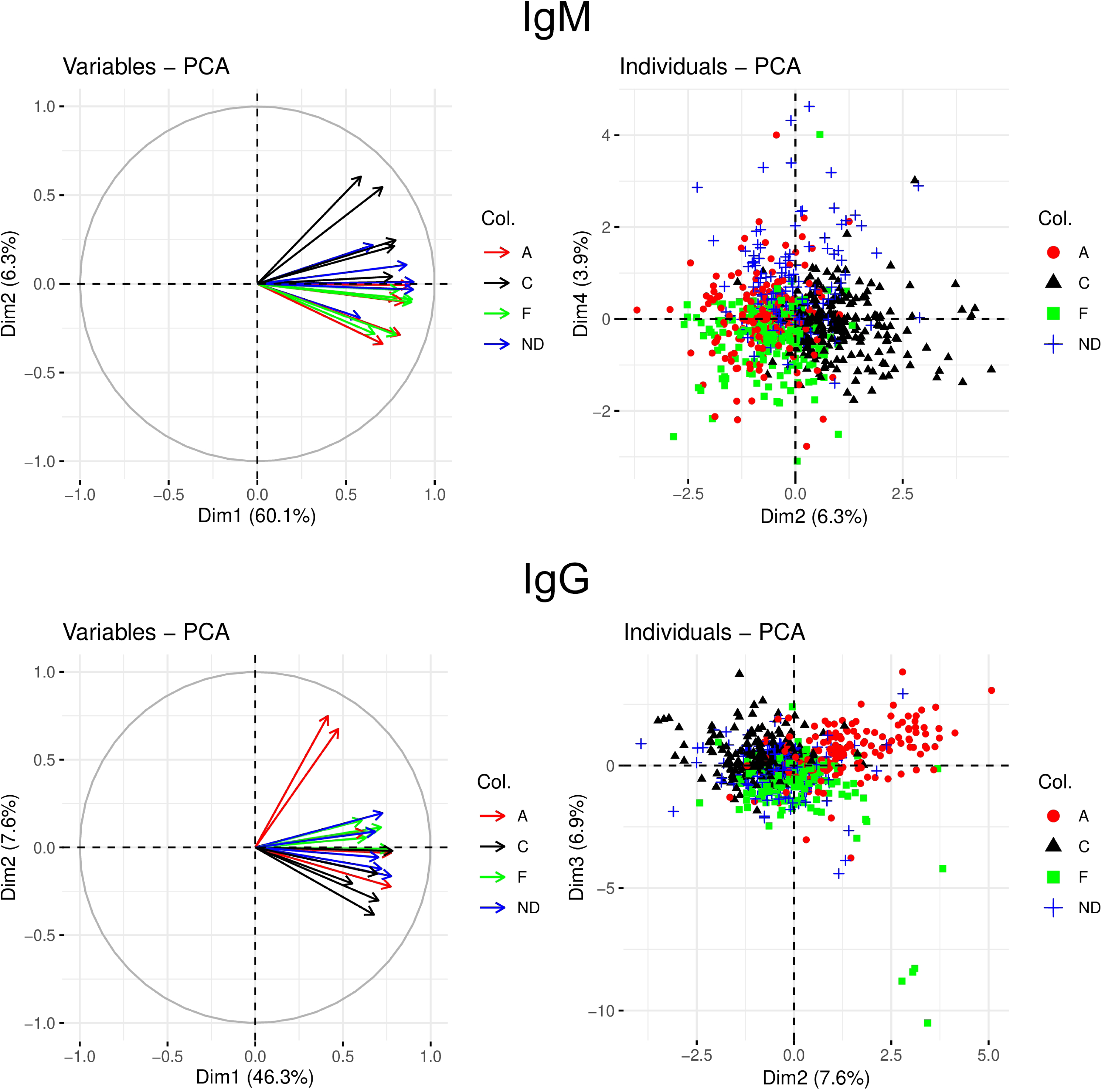
Principal component analysis (PCA) of the binding data of IgM (top) and IgG (bottom) from patients with AD (red), FTD (green), NDD (blue), and healthy controls (black) to an array of 722 7-mer mimotopes of human public IgM reactivities. The PCA analysis was used to characterize the reactivities by their patterns across patients, so the layout is transposed, and “individuals” refers to individual reactivities. The variables are the respective serum samples. Left – correlation of serum reactivities mapped to the first two components – the first component correlates highly with all sera and accounts for 60% of the variation. The raw data have been normalized for the physicochemical properties of the amino acid residues in the peptides. Right – individual mimotope reactivities color coded for the four diagnostic groups studied, mapped to factors 2 and 4 (IgM) and 2 and 3 (IgG).

### 3.2. Graph representation of mimotope reactivities for a detailed analysis

The PCA, indicated that the public IgM reactivity repertoire contains information relevant to the neurodegenerative disease under study. To extract further detail, the cross-reactivity data for both isotypes were represented as graphs along the lines of the methodology published recently (Ferdinandov et al., 2023; Pashova-Dimova et al., 2023). The method for inferring cross-reactivities between peptides was improved by measuring the similarities between the profiles observed in the set of sera (see Materials and methods for a detailed description). The threshold of profile similarity for connecting two mimotope reactivities with an edge was adjusted to yield a large connected component of the graph with at least 700/722 reactivities. The connected components of the graphs were visualized in 2D using a two-step spectral embedding (Suppl. Fig. 2). A 3D final projection is also provided, which is of even higher fidelity to the four or 5D spectral embedding (Supplementary files IgM_AD_FTD_NDD_C.mp4 and IgG_AD_FTD_NDD_C.mp4). The visualizations illustrate a very clear distinction between the public reactivities lost (high in the controls) and those gained in the patients with dementia. Furthermore, there is also a clear distinction between the diagnoses, especially in the IgG graph.

### 3.3. Reactivity graph structure and parameters

To estimate the significance of some descriptive parameters of the graphs, the baseline values were simulated by using graphs generated by the same algorithm but based on the scrambled raw data matrices for each isotype. The graph density (the number of edges relative to the maximal number of possible edges) and the size of the large connected component were higher for both the IgM and the IgG graphs then for the random graphs (Fig. 3). Similarly, the degree distributions were shifted slightly to the right (Suppl. Fig. 3) and the maximal clique size distributions showed more cliques but with similar distribution as compared to the random graphs (Suppl. Fig. 3).

**Figure 3.**
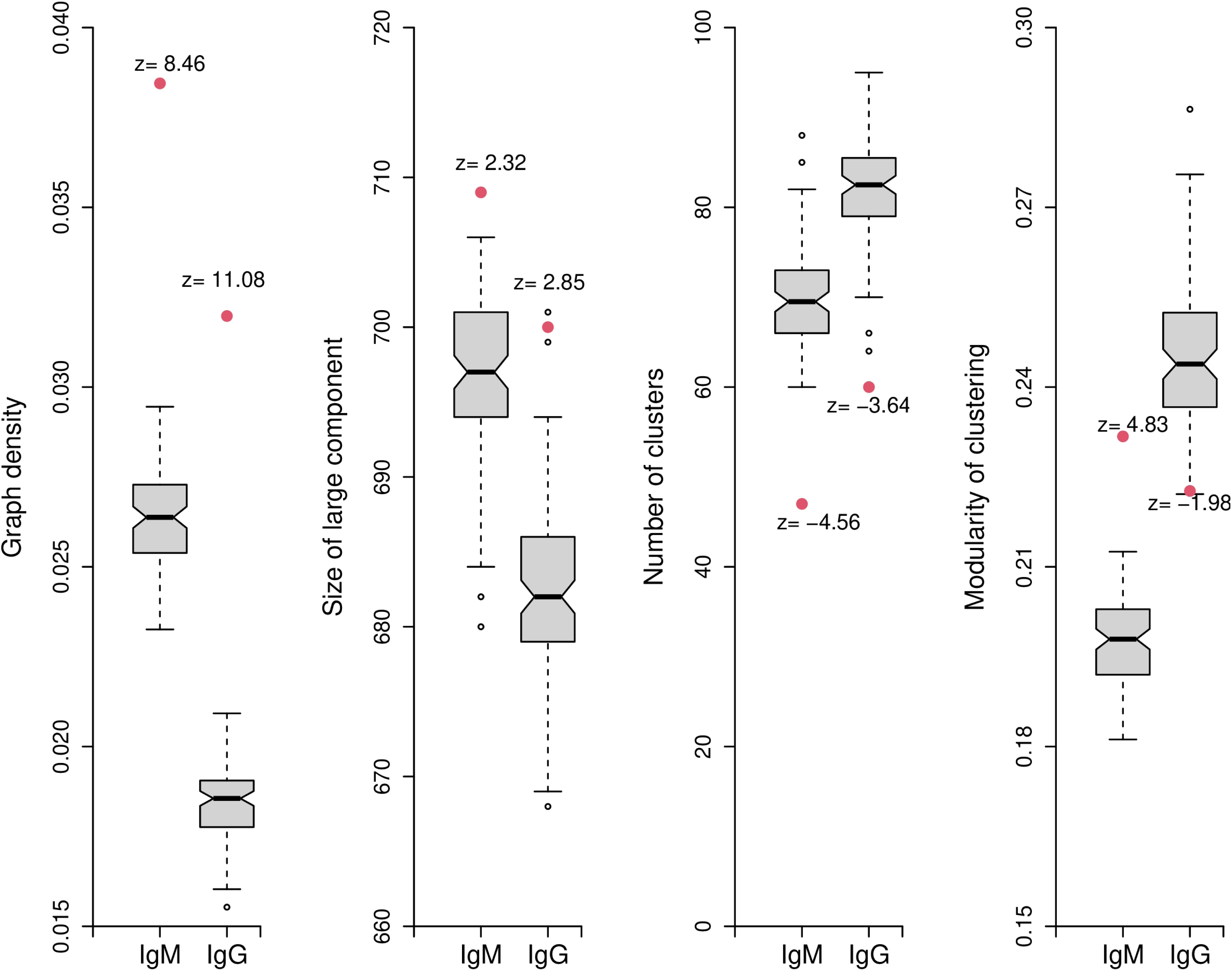
Some parameters of the IgM and IgG reactivity graphs. Each parameter has been bootstrapped by generating 100 graphs constructed using the same algorithm but using scrambled matrices of the raw data. The antibody reactivity graphs are dense and clustered into fewer clusters than the random graphs. Still, the modularity of the clustering is very different for IgM (higher) and IgG (lower) than that of the random graphs.

Next, the information about classes of public reactivities encoded in the topology of the graphs and their relation to neurodegenerative diseases was studied. Clusters of vertices were determined in the graphs based on modularity (regions with denser edges within than to other parts of the graph) using the Leiden algorithm (https://github.com/TomKellyGenetics/leiden). The IgM graph had 47 clusters, with 13 consisting of single vertices and the largest containing 60 vertices. For the IgG graph, these values were 60, 22, and 33, respectively. The IgG graph clusters appeared more numerous and smaller in size, but these differences were not significant. Compared to the random graphs, both the IgM and the IgG graphs had a significantly lower number of clusters (Fig. 3), indicating that the repertoire of public reactivities at this coarse resolution exhibits large-scale modularity with relatively few clusters. Interestingly, compared to the random graphs, the modularity of the clustering was very different between IgM (high) and IgG (low), although the values for both graphs were rather similar. This can be interpreted as the public IgM repertoire clustering into a relatively small number of tight clusters, while the IgG reactivities to the IgM mimotopes form slightly more clusters that are significantly less well-defined.

The vertices of the graphs were assigned attributes corresponding to the rank sums of each mimotope’s reactivities in different diagnoses. Thus, for instance, mimotopes with high reactivities in the control and low in the disease groups will have a high rank sum associated with the control group. The attributes also reflected blood group antigen cross-reactivity, as well as cross-labeling – assigning the IgM graph vertices the IgG attributes by diagnosis and vice versa. The latter approach was applied to check if the graph of each isotype reflects the diagnosis-associated relations of the other.

The degree of correlation of the graph structure with a quantitative attribute is measured by assortativity – the extent to which neighboring vertices have similar magnitudes of the attribute. Figure 4 compares the assortativities of several characteristics for both graphs compared to the respective sets of random graphs. Here, assortativity was intended to indicate the degree to which reactivities, correlating with the respective property, cluster. The highest assortativity was found for the control group-related reactivities (i.e., those lost in disease). The assortativity of the correlation with blood group antigens was comparable but somewhat lower, and the cross-labeling attribute assortativities were markedly lower, reaching insignificant levels in FTD (for IgM) and NDD (for IgG). The correlation between the reactivity profiles across isotypes (GMCR) did not show significant assortativity. Thus, the clusters of disease-specific cross-reactivities among these public IgM mimotopes did not match.

**Figure 4.**
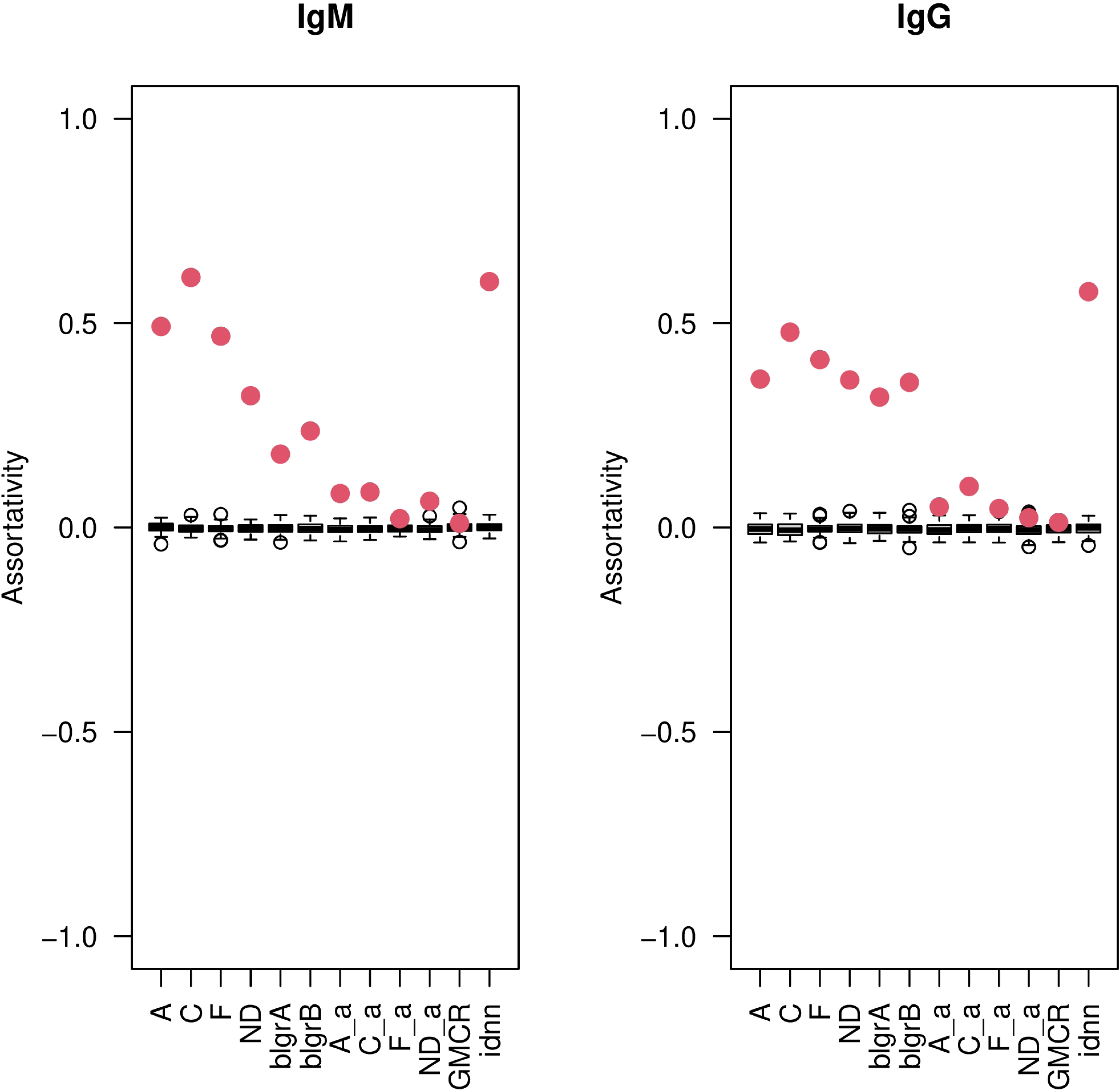
Assortativity of the properties associated with the vertices (mimotopes) for the IgM (left) and the IgG (right) reactivity graphs. The assortativity measures the degree to which vertices with similar values are connected; respectively, a highly assortative property segregates the mimotopes in the space of their reactivities. The values for the studied sera are compared to the values obtained for random graphs (100 graphs constructed using the same algorithm but using scrambled matrices of the raw data) and using the same labeling of the vertices. The highest assortativity is observed for the control samples (C), as well as for the AD (A) and FTD (F). Cross-labeling the vertices using the reactivity magnitude seen in IgM for the IgG graph and vice versa gives much lower assortativity.

### 3.4. Idiotypy and neurodegenerative disease

In our previous studies, we stumbled upon a significantly high proportion of homologies to HCDR3 region sequences from human antibodies in the public IgM reactivity mimotopes (Pashov et al., 2019; Pashova et al., 2022). Such HCDR3 homologues of mimotopes could be considered, at least potentially, as idiotopes. Since the IgM mimotope array used in this study was shown previously to be significantly enriched in idiotipic mimotopes, we decided to check if this idiotope reactivity correlated with reactivity associated with neurodegenerative diseases. The nearest neighbors for each mimotope among a library of 4×10^6^ 7-residue frames from 10^6^ human antibodies were determined using a criterion of at most one mismatch. Under this condition, 483/722 mimotopes had at least one nearest neighbor among the J region frames, and some had up to 187. Probably, the latter coincide with germline genes. Next, the number of nearest neighbors among idiotopes (NNNI) that best splits the data into groups correlated with diagnosis was determined. To that end, the mimotope set was repeatedly divided into two groups relative to each value of the sorted NNNI. The correlation between mean reactivity rank by disease and the grouping of NNNI low and high mimotopes was calculated using the Wilcoxon rank sum test at each step. Figure 5 shows that for IgM, the finding of 2 idiotopes among the nearest neighbors was the optimal criterion to detect maximal correlation. The correlation was positive for the reactivities lost in the disease groups (high in the control group) and negative for the reactivities high in AD. The group of ND also had a positive correlation, although much lower than the group C, and the FTD group had a negative correlation similar to AD, but much lower. For IgG, these correlations were mainly insignificant. Unexpectedly and contrary to the IgM, the IgG reactivities lost in neurodegeneration (C) had a small but significant negative correlation. In contrast, the ND group had a slight positive correlation, like in AD. Thus, the public IgM reactivities lost in the neurodegenerative diseases tend to be more idiotypically connected, in contrast to, in particular, those gained in AD but also to some extent those found in FTD.

**Figure 5.**
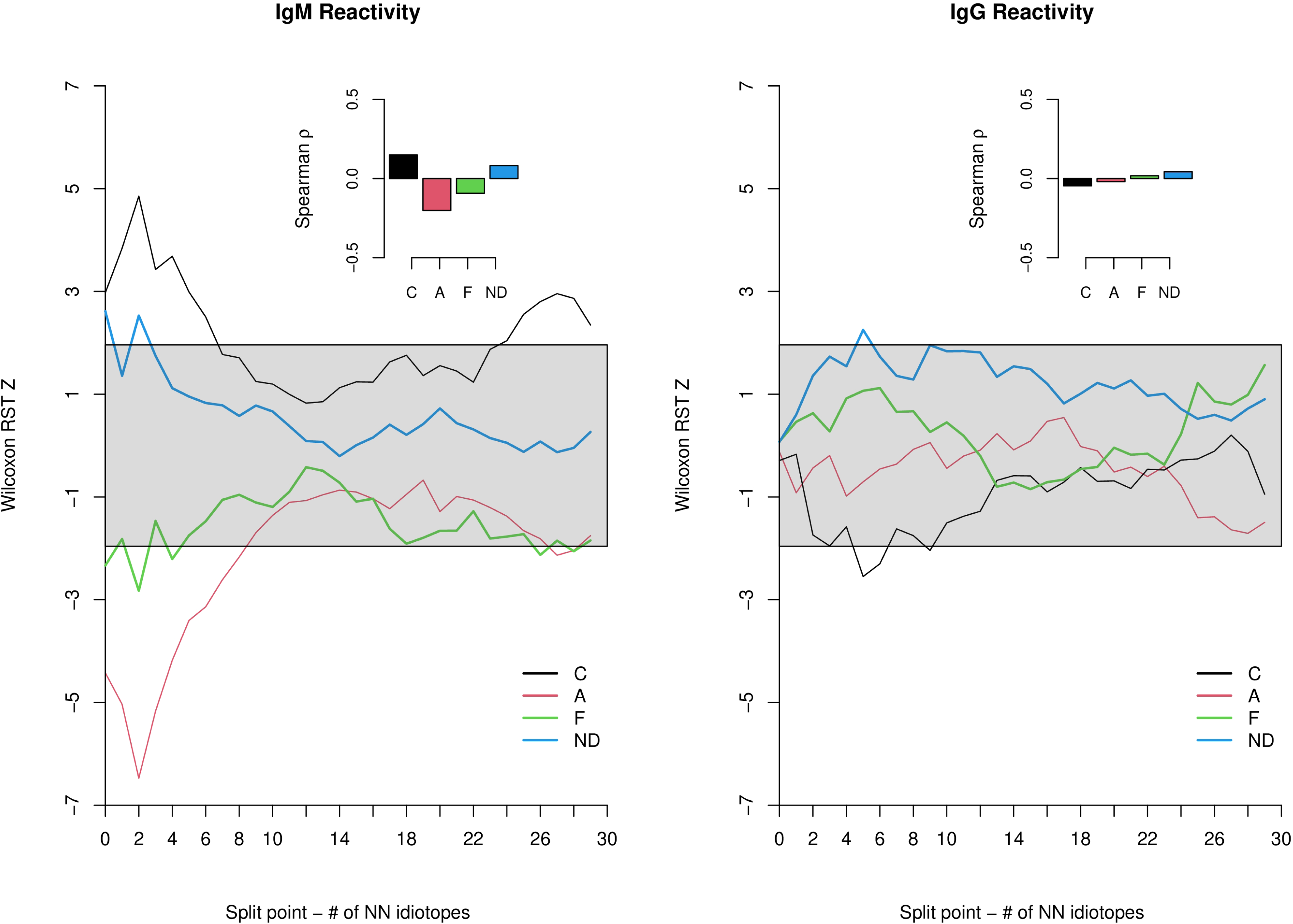
Correlations between IgM and IgG reactivity averaged by diagnosis and number of nearest neighbors among idiotopes. Red – AD (A), green – FTD (F), blue – NDD (ND), and black – controls (C). Wilcoxon rank sum test was used to compare the reactivities in the peptides dichotomously grouped by splitting the mimotope set relative to the number of neighboring idiotopes of each peptide. The test was performed at each split point indicated on the abscissa. The resulting z-score is on the ordinate. The shaded area indicates a significance level p≥0.05, so the portions of the curves outside of the shaded area correspond to a significant difference. The insets show Spearman’s correlation coefficient for the reactivity vs the number of idiotope neighbors. The IgM reactivities, higher in the controls (i.e., lost in the other patients), show the highest positive correlation with idiotypy, while those increased in AD show the highest negative correlation.

### 3.5. Public reactivity clusters associated with neurodegenerative diseases

The mean reactivities by diagnosis in each vertex cluster, defined in 3.3, were compared using the Dunn test (BH - Benjamini & Hochberg adjusted p<0.01). The clusters found to have significant differences between at least two diagnoses were 17/47 for the IgM graph and 17/60 for the IgG one. To summarize these results, each significant cluster was represented by the six differences between mean reactivities for the four diagnoses. The data was subjected to PCA and plotted as a biplot (Fig.6), which shows simultaneously the correlations between the axes as well as the mapping of the clusters onto the same space. The IgM graph clusters show differences mostly along the axis defined by differences between the controls vs AD and FTD, with a weak correlation between the control reactivities and the ND patients. The IgG clusters complemented these distinctions by differences between AD and FTD. The symmetric distribution of the clusters around the origin shows that there are approximately equal numbers of clusters representing positive or negative differences.

**Figure 6.**
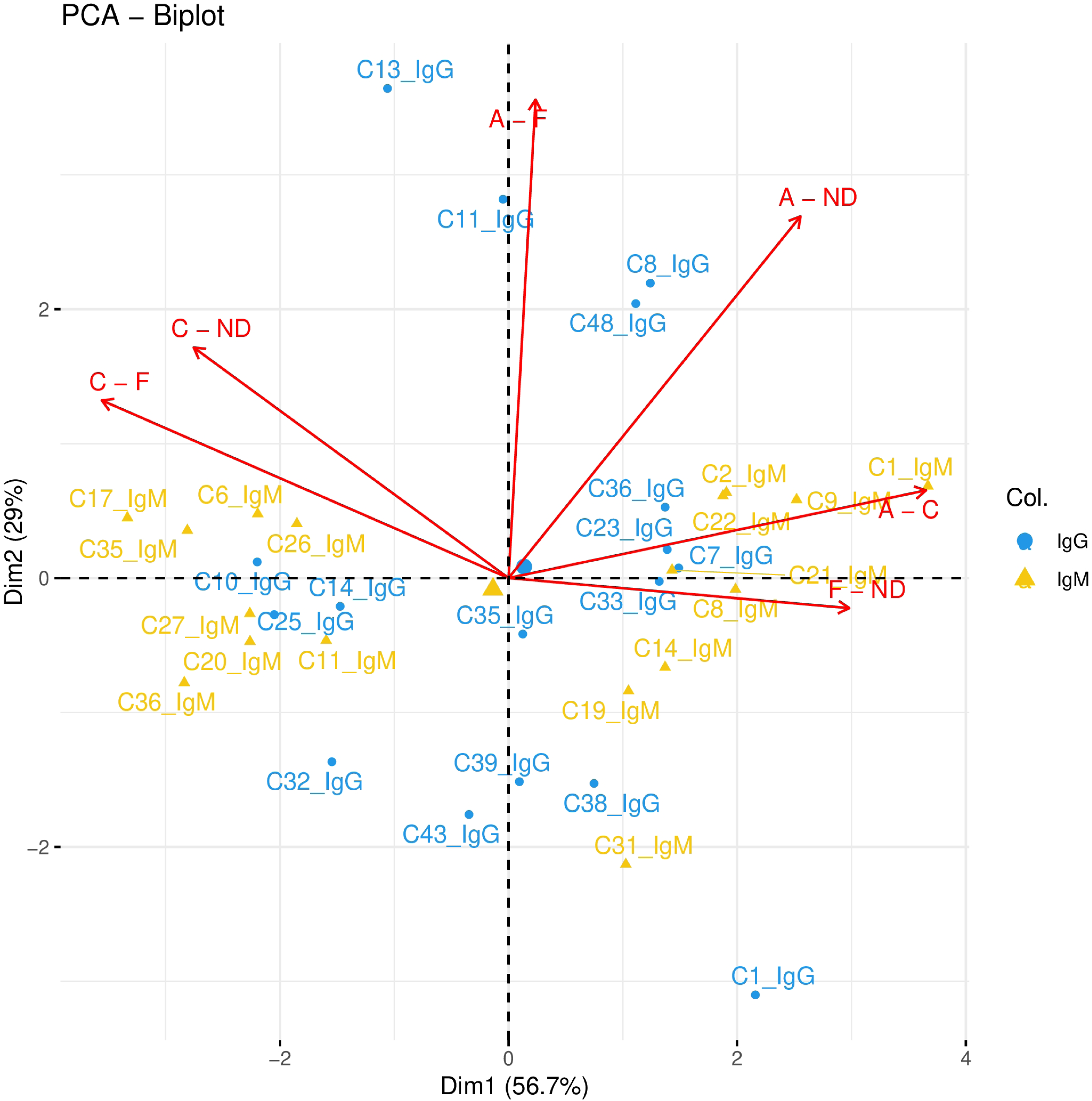
Biplot from the PCA results on the differences between the mean reactivities by diagnosis in the established clusters in the IgM and the IgG graphs. Only the clusters for which the differences are significant at the level p<0.05 after adjusting for false discovery rate using the Benjamini-Hochberg correction are used. IgM shows a simpler separation of the clusters, mainly between those with high reactivity in AD and FTD and those with low reactivity in the patients (high in the controls). The IgG clusters show higher diversity, some differentiating AD and FTD or NDD, but their differences were generally smaller than those in the IgM clusters.

The repertoire probing with just 722 mimotopes provides too coarse a picture. The reactivity graphs clustering summarized the cross-reactivities based on profile similarities. The respective underlying reactivities are encoded in the pattern of similarities between the samples that generate an edge. So, although the reactivities in each cluster were generally related, augmenting the grouping by a further clustering of the edges would provide a higher resolution. Indeed, when each cluster was subdivided into (overlapping) subclusters connected by highly correlated edges, a new, more detailed clustering was obtained. It consisted of 211 clusters (sizes 3-47) for the IgM graph and 236 clusters (sizes 3-26) for the IgG graph. Applying similar strategy for statistical analysis as for the vertex clusters with a significance level of p<1e-3 after BH adjustment, 44 highly significant secondary clusters were found for the IgM graph and 21 – for IgG (Fig. 7). At an even higher level of significance (p<1e-5) 18 IgM clusters and 11 IgG clusters remained. Many of these clusters had the same type of difference between the diagnoses, so they could be grouped into seven groups for the IgM and four for the IgG (Suppl. Table 2). As concluded above, IgM reactivities did not differentiate between AD and FTD, but IgG clusters did. Since the final clusters of mimotopes were overlapping, their relations were examined using a Venn diagram (Suppl. Figure 5). The clusters with higher reactivity in the controls were only from the IgM set, while those with disease-associated reactivities tended to overlap between IgM and IgG. The IgG reactivities, which were high in disease, grouped in more diverse clusters. Interestingly, 2 clusters were high in the controls and low in FTD in IgG and vice versa for IgM.

**Figure 7.**
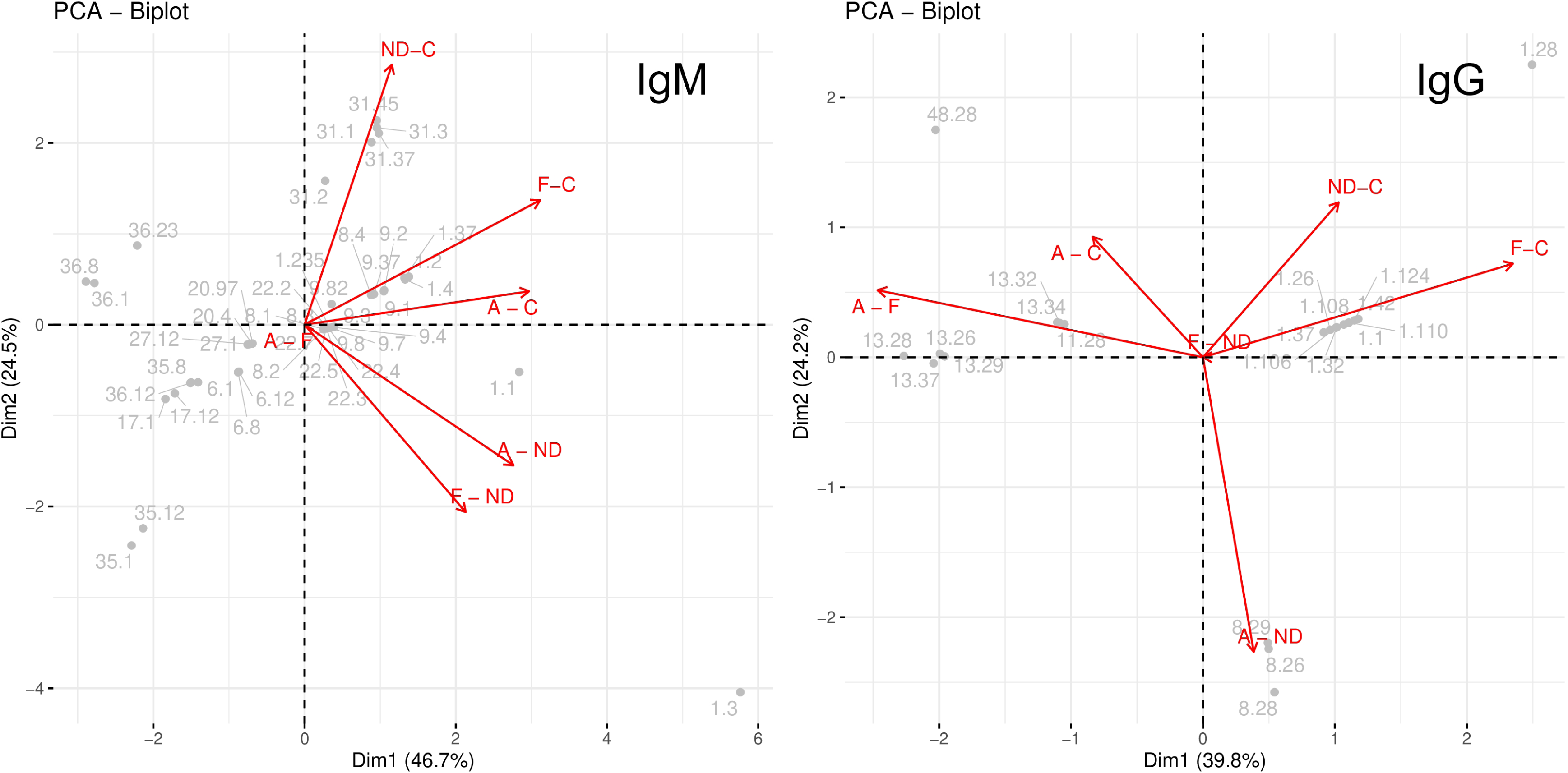
Biplot from the PCA results on the differences between the mean reactivities by diagnosis in the established combined (vertex and edge) clusters in the IgM and the IgG graphs. The statistical parameters are the same as for Fig.7. Again, no IgM cluster differentiates AD and FTD, while IgG clusters do. The cluster numbers comprise the vertex cluster separated from the subdividing edge cluster by a point.

### 3.6. Mapping the NNNI to clusters

Since the disease associated reactivities were clearly correlated to idiotypy, at least in the IgM data set, the mean NNNI of each of the final clusters were compared (Fig.8). As expected, for IgM the clusters of reactivities higher in AD and FTD had lower NNNI then those lost in the dementia patients (C) or the overall mean for the remaining (not significant) clusters (labeled ALL). Among the reactivities lost in dementia as compared to the controls (high in C), only those higher in ND than in C had significantly more idiotope neighbors than the bulk (ALL). In IgG, surprisingly, only the clusters with reactivity higher in FTD than in the controls had significantly higher NNNI.

**Figure 8.**
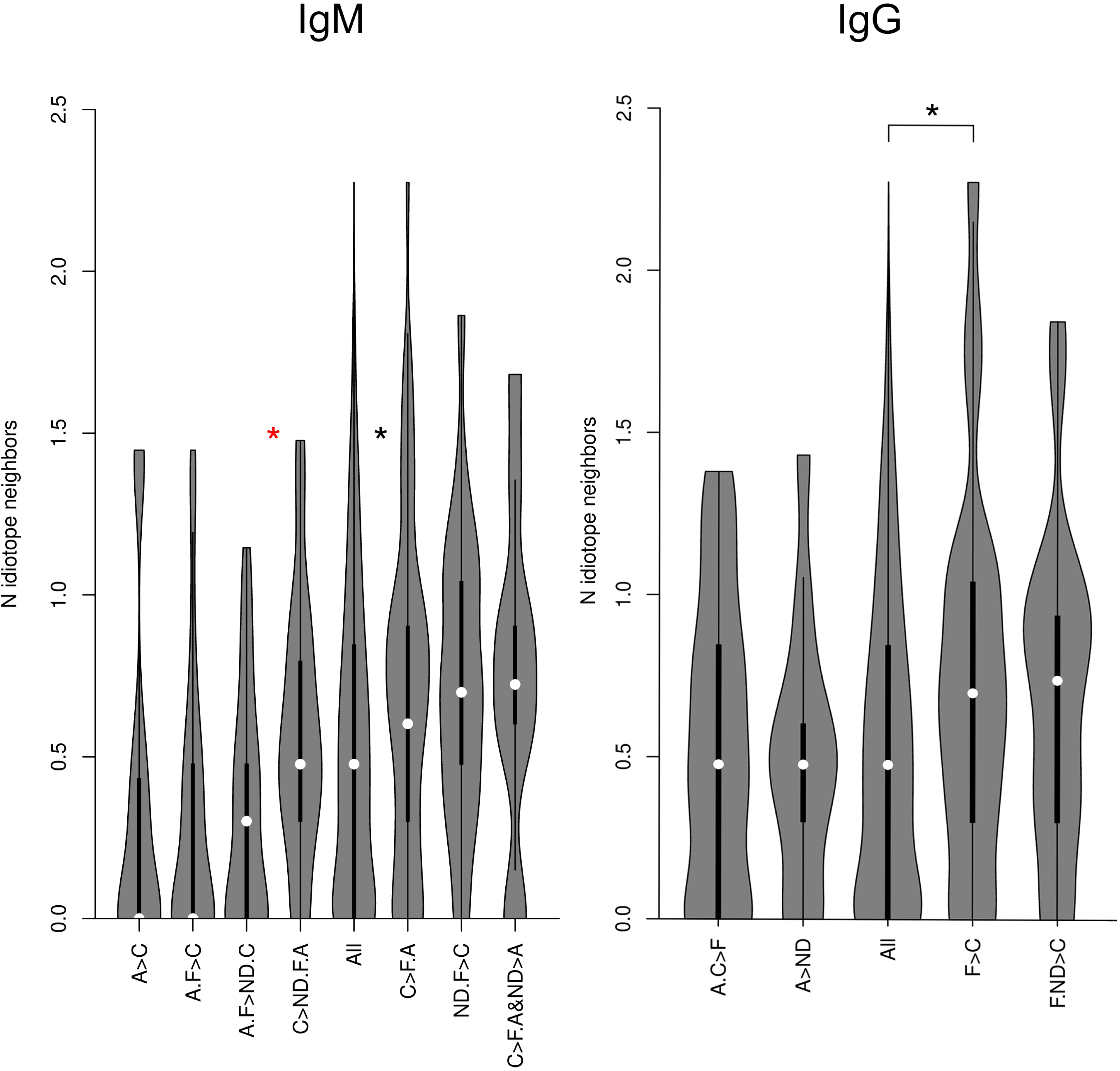
Distribution of the decimal logarithms of the number of idiotope neighbors by cluster groups. Since many significant clusters had the same reactivity pattern across diagnoses, they were reunited to form cluster groups with the same difference in reactivity between diagnoses. These cluster groups are analyzed to determine the distribution of the number of idiotope neighbors per peptide. For comparison, the set of tested mimotopes not included in the selected clusters was added as a separate group (All). The groups are sorted by the increasing mean number of idiotope neighbors. Dunn’s test showed that, for the IgM graph, the clusters to the left of the red asterisk have fewer idiotope neighbors than the rest. Also, the groups with higher expression in the controls as compared to AD and FTD (C>F.A) had more idiotope neighbors than the mean for those not included in the significant clusters (black asterisk). For IgG, only the reactivities in FTD had more idiotope neighbors than those not included in the significant clusters.

### 3.7. Mapping the altered reactivity mimotopes to potential self-antigen linear epitopes

The reactivity of public antibodies, especially IgM, could often be identified as directed to carbohydrate epitopes on bacteria. Still, many of these reactivities probably come from natural antibodies which have no nominal antigen in the sense of adaptively evolved specificity and are inherently poly/autospecific. Also, it is almost impossible to deconvolute conformational protein epitopes corresponding to mimotopes selected by a bulk repertoire. With these considerations, the only accessible specificity interpretation of the studied mimotopes was to check them against potential linear epitopes in the human proteome. BLAST search for sequences which had at most one mismatch with any of the studied mimotopes yielded 184/722 sequences with hits in 2182 proteins and 46/197 sequences with hits in 67 proteins for the mimotopes selected in the ultimate disease-associated clusters (Suppl. Table 3). Among the 67 putative targets, 31 were involved in brain metabolism and signaling. The proteins LRPs (LRP1B, LRP4, LRP6, and proLRP), sortilin, TIA-1, and BMAL1 are directly contributing to the pathogenesis of neurodegenerative disorders. The LRPs are endocytic receptors for the internalization of ApoE-containing lipoproteins, and sortilin is an endocytic receptor for the uptake of diverse proteins, including ApoE (Asaro et al., 2021; Carlo et al., 2013), amyloid beta (Saadipour et al., 2013), and α-synuclein (Ishiyama et al., 2023). These receptors had homologies with mimotopes that were part of a reactivity group overexpressed in IgG from FTD and ND patients. TIA-1 - a cytoplasmic scaffold protein of stress granules, was homologous to mimotopes of reactivities overexpressed in IgM from FTD and ND patients. BMAL1 – a transcription factor which is an important regulator of circadian rhythm, has a mimotope in a group of reactivities overexpressed in IgM from AD and FTD. Some other potentially targeted proteins with involvement in neurodegeneration are SMG7 (mRNA quality control), Tenascin C (neuroinflammation and association with core amyloid-beta plaques), FGF17 (cognitive processes), IRS2 (Insulin receptor signaling), Lon protease (protein quality control), UBXN1 (protein quality control), ZFYVE26 (autophagy), PlexinD1 (neuronal guidance) (Suppl. Table 4).

### 3.8. Generalizability

To assess the separability and generalizability of the observed patterns, a cross-validation scheme, leaving out sets of pools with no common serum with the rest of the samples, was applied to an SVM model. The Materials and Methods section gives details on the feature selection and cross-validation. The models were trained separately on the IgM and the IgG data and a joined data set of IgM and IgG reactivities. The only dichotomy that yielded significantly high accuracy (0.85) in the cross-validation scheme was AD/FTD vs C/ND in the IgM data set.

## 4. Discussion

The presented study aimed at exploring the capacity of the public IgM antibody repertoire to detect and react to different types of dementia. To this end, the public IgM repertoire and the IgG counterparts that are cross-reactive with it were analyzed using high-throughput binding assays and cross-reactivity graphs. The immune system’s role in neurodegenerative diseases has been supported by extensive evidence, both in terms of cellular immunity (Chen et al., 2018; Reagin and Funk, 2023; Yang et al., 2022), inflammatory mechanisms (Baulch et al., 2020; Foley et al., 2024; Shue et al., 2024), and antibody-based biomarkers (DeMarshall et al., 2022; DeMarshall et al., 2023; Kheirkhah et al., 2020; Restrepo et al., 2013). Particularly, concerning the antibody repertoire, several groups have demonstrated the utility of IgG adaptive immunity responses as a source of biomarkers, including for early detection (DeMarshall et al., 2016; Reddy et al., 2011; Restrepo et al., 2013). The primary paradigm underlying these studies was basically autoimmunity, possibly a “protective” one (Yoles et al., 2001). At the same time, others considered the role of constitutively expressed natural antibodies (Rothstein, 2016), although in most cases, the term naturally occurring is used for antibodies elicited by self-antigens (Britschgi et al., 2009; Rosenmann et al., 2006; Szabo et al., 2008).

Enticed by the successful mining of the antibody repertoires for AD biomarkers (DeMarshall et al., 2015; DeMarshall et al., 2016; Restrepo et al., 2013), we looked for public IgM reactivities with the same properties. A recently developed approach was applied (Ferdinandov et al., 2023; Pashov et al., 2019; Pashova et al., 2022). It consists of combining IgOme repertoire analysis (Ryvkin et al., 2012) with the immunosignature concept (Restrepo et al., 2013). The sera were studied in pools of two to emphasize the public reactivities and reduce the cost of the microarrays. The small number of samples had already precluded the applicability of applicable machine learning models.

Compared to controls, we could demonstrate distinct patterns of public IgM and their cross-reactive IgG counterparts in AD, FTD, and ND patients. The loss or gain of specific IgM reactivities in AD and FTD indicates alterations in the natural and possibly the induced IgM repertoire. The loss of constitutive natural antibody reactivities has been observed before (Ferdinandov et al., 2023; Pashova et al., 2022). It probably represents a temporary exhaustion due to binding to some overexpressed self-antigens in terms of a scavenging function for cellular debris, as well as quantitative tolerance thresholds (Hennings et al., 2011). The opposite, reactivities increased in disease, is most easily explained as an adaptive immune response.

The concept of public clones has two aspects. Antibodies can be shared between individuals as clonotypes or just as a convergent specificity. In the latter, broader context, the IgM repertoire is not only more diverse but also organized in larger public specificity classes compared to IgG or IgA (Arora and Arnaout, 2022; Galson et al., 2015; Wu et al., 2022). It is also safe to assume that natural antibodies and public IgM overlap without being synonymous. Sometimes, the IgM compartment is considered to consist predominantly of ill-defined polyreactivities or even antibodies that are simply sticky (Ausserwöger et al., 2022; Cunningham et al., 2021). Previously, graphs were used to analyze the repertoire complexity (Miho et al., 2019) and antibody cross-reactivity patterns (Ferdinandov et al., 2023; Madi et al., 2011). The cross-reactivity graph provided a framework for visualizing and analyzing the IgM network of specificities inferred only from high-throughput binding assays. Spectral embedding ensured a well-defined visualization that can provide insight into the graph topology.

Our study found the cross-reactivity graphs to be dense and clustered in a relatively small number of clusters. It is possible that this is the image of the classes of overlapping reactivities discussed by Arora and Arnout (Arora and Arnaout, 2022). It is seen at a relatively coarse resolution due to the low diversity of the mimotope library relative to the antibody diversity. The clusters had a higher modularity in IgM, indicating that their structure is less distinct in the IgG graph, probably due to addressing IgM mimotopes with IgG. Thus, the IgG reactivities were either randomly cross-reactive, IgG antibodies, or isotype-switched and probably mutated counterparts of the observed IgM. The characteristic large-scale cross-reactivity patterns of AD and FTD were manifested as high assortativity—neighboring mimotopes had similar levels of reactivity in a given diagnosis. The highest assortativity was observed for the control group, emphasizing the loss of IgM reactivities common to the different neurodegenerative diseases. Since the edges were defined as binding pattern correlations across samples, high assortativity indicates the prevalence of cross-reactivities between mimotopes for which the measured property is similar. Cross-reactivity groups can also be interpreted as footprints of major antibody reactivities with the respective properties. Like all other cross-reactivity graph observables, assortativity depends on the set of probes used. By sampling symmetrically the sequence space of a library of public IgM mimotopes, the probe set is optimized simultaneously for diversity and cross-reactivity (Pashov et al., 2019). Our results also show that it extracts information from the IgG repertoire with similar efficiency but different patterns, pointing to the relations between the public IgM repertoire and the induced IgG repertoire.

The observations from the assortativity analysis were studied in detail by the distribution pattern of these properties among the graphs’ clusters. In the IgM network, the key distinction was between the controls (correlating with ND) vs AD and FTD. The latter were not well separated in the clusters. The good distinction between AD and FTD was apparent only in the IgG data set. Interestingly, FTD-associated IgG not only differed from those in AD but, unlike IgM, they were correlated with ND. Unlike AD, FTD IgG patterns appeared to have higher idiotypic connectivity. These facts point to the occurrence of new IgG reactivities in FTD, which are mostly unrelated to the public IgM repertoire. The distinction of ND from AD and FTD, and its similarity at times to the controls, could be characteristic of just some of the diagnoses in this heterogeneous group.

The BLAST search of the human proteome produced hits of proteins, potential targets of the reactivities found associated with dementia. Many of those putative autoantigens are implicated in neurodegeneration. This is especially true for the endocytic receptors LRPs and sortilin, which, although very different in structure, share ApoE and ApoE/amyloid beta complexes as ligands for internalization and subsequent endolysosomal degradation (Cam et al., 2004; Chow et al., 2021; Kanekiyo and Bu, 2014; Zhang et al., 2020) (Asaro et al., 2020). The clearance of amyloid beta from the brain was shown to depend on the level of expression of LRPs (Castellano et al., 2012), and sortilin-deficient mice present elevated levels of APOE and increased amyloid beta accumulation (Carlo et al., 2013). Additionally, LRP4 and LRP6 are engaged in Wnt/beta-catenin signaling, an important pathway promoting cell proliferation during early development, which is tightly controlled during aging and suppressed in neurodegeneration (Inestrosa and Toledo, 2008). Antibodies to these LRPs may contribute to the pathology of neurodegeneration by causing a loss of control over the Wnt/beta-catenin pathway. A tau-related neurodegenerative factor - TIA-1 was also identified as a target. It is an RNA-binding protein and a scaffold for the formation of stress granules (SG) in response to cellular stress. Downregulation of TIA-1 restrains the inclusion of tau in pathological and persistent stress granules, while overexpression of TIA1 stimulates tau-related degeneration (Apicco et al., 2018; Maziuk et al., 2018). Another target, a factor etiologically related to neurodegeneration and associated with sleep disturbances in AD, is the circadian transcription factor BMAL1. It is the main driver of circadian rhythm and is involved in the rhythmic transcription of about 5 to 10 % of expressed genes (Hughes et al., 2009). BMAL1 has been shown to be aberrantly methylated and irregularly expressed in the brains of AD patients (Cronin et al., 2017; Hulme et al., 2020).

A number of previously discussed biomarkers for FTD and AD, like Nfl, progranulin, TDP-43, or GFA (Duran-Aniotz et al., 2021) were not among the potential targets of the antibody reactivities observed, most likely due to the coarse repertoire probing. Notably, a recent study found 7 specific serum autoantibodies to MAPT, DNAJC8, KDM4D, SERF1A, CDKN1A, AGER, and ASXL1 and proposed a classification model that differentiated AD from other neurodegenerative diseases with high accuracy (Fang et al., 2024). Furthermore, using proteome-wide autoantibody screening, Matsuda et al. identified 229 antibodies differentially elevated in AD or dementia with Lewy bodies (Matsuda et al., 2025). Autoantibodies targeting neuropeptide B and adhesion G protein-coupled receptor F5 showed significant correlations with clinical traits.

Previously, we found that the public IgM mimotope library is enriched in homologues to HCDR3 sequences of human immunoglobulins. Looking into this correlation, the public IgM reactivities lost in AD and FTD proved significantly cross-reactive with idiotopes. On the contrary, the IgM reactivities increased in AD (and partially those increased in FTD) were much less idiotypically connected. The correlation between idiotypic connectivity and disease-specific reactivities points to a potential mechanism of an AD-associated immune dysregulation, probably shaped by the induced autoimmunity. Interestingly, DeMarshall, et al. defined a diagnostic autoantibody reactivity panel for AD, including auto-light chain kappa antibodies (DeMarshall et al., 2023). Another interesting aspect is revealed concerning the recent finding on the pivotal role of the peripheral immune system in the brain-gut-periphery axis, potentially shaping the trajectory of AD (Han et al., 2018; Iban-Arias et al., 2024). The intriguing link necessitating further studies is through the role of the microbiome in shaping the IgM repertoire (Li et al., 2020) and the possibility that the idiotypic connectivity is at least partially, if not mainly, determined during repertoire selection (Detours et al., 1996; Kazatchkine et al., 1994).

Only 9/65 different proteins had putative linear epitopes differentially bound by IgM in the controls (reactivities disappearing in disease), in contrast to the prevalence of idiotypic connections in IgM compared to IgG. The mimotope sequence YWTDSSR has different one-residue-mismatched homologues in LRP1 B, LRP4, and LRP6 (see supplemental table 3). It represents the beta-propeller domain signature, found in most LRP, and is characterized by high polyspecificity to multiple ligands (Príncipe et al., 2021). A NCBI protein blast shows that the exact mimotope sequence is highly conserved across many species of bacteria and fungi. Furthermore, YWTDSSR proved to have seven idiotope neighbors in our analysis. Finally, it was found in the IgG cluster of reactivities increased in FTD and ND. Thus, at least some patients in the heterogeneous ND group showed similarity to FTD concerning this autoreactivity. So far, anti-LRP autoantibodies have been associated only with myasthenia gravis (Chuquisana et al., 2024). This interesting example highlights the connection between autoantibodies associated with neurodegeneration, antibody polyspecificity and other promiscuous protein binding patterns, public IgM, and HCDR3 idiotopes. Taken together, all these aspects of autoimmunity in neurodegenerative diseases are possibly related by processes involved in repertoire selection.

Thus, the differences in antibody public IgM reactivity profiles highlight another repertoire compartment with considerable potential as a source of non-invasive biomarkers for early disease diagnosis. Despite the promising findings, this study has several limitations. The experimental setting with small groups and overlapping serum pools precluded important model validation, so only a rudimentary classifier was trained to prove separability, at least of AD/FT vs C/ND. Furthermore, the overlapping pooling requires caution in interpreting correlations. Thus, the conclusions should be considered preliminary. Nevertheless, our results also clearly indicate the utility of the reactivity graph approach and the significance of the association of idiotypically connected IgM reactivities with the immune pathogenesis of neurodegenerative diseases.

Additionally, the study primarily focused on linear mimotopes. Conformational epitope information is encoded in the clusters of linear mimotopes, but is hard to deconvolute without knowing in advance at least the target antigen, if not the exact epitope. This task remains to be addressed in future studies.

In conclusion, this study provides proof for the utility of public, idiotypically connected IgM reactivity profiles as biomarkers for AD and FTD. Applying cross-reactivity graphs to high-throughput antibody binding data offers a powerful tool for uncovering complex patterns and networks associated with disease states.

## Supporting information

GraphEmbeddinh_3Danimation

## Declaration of generative AI and AI-assisted technologies in the writing process

During the preparation of this work, the author(s) used Grammarly to improve the style of the manuscript. After using this tool/service, the author(s) reviewed and edited the content as needed and took full responsibility for the content of the publication.

## 5. Acknowledgements

This work is supported by grants # KP 06 N 33/5/2019 and # KP 06 N 37/24/2019 from the Bulgarian National Science Fund. The Bulgarian cohort was supported by the European Union-NextGeneration EU (National Recovery and Resilience Plan of the Republic of Bulgaria, grant number BG-RRP-2.004-0004-C01).

## Supplementary Materials

**Supplementary Table 1.**
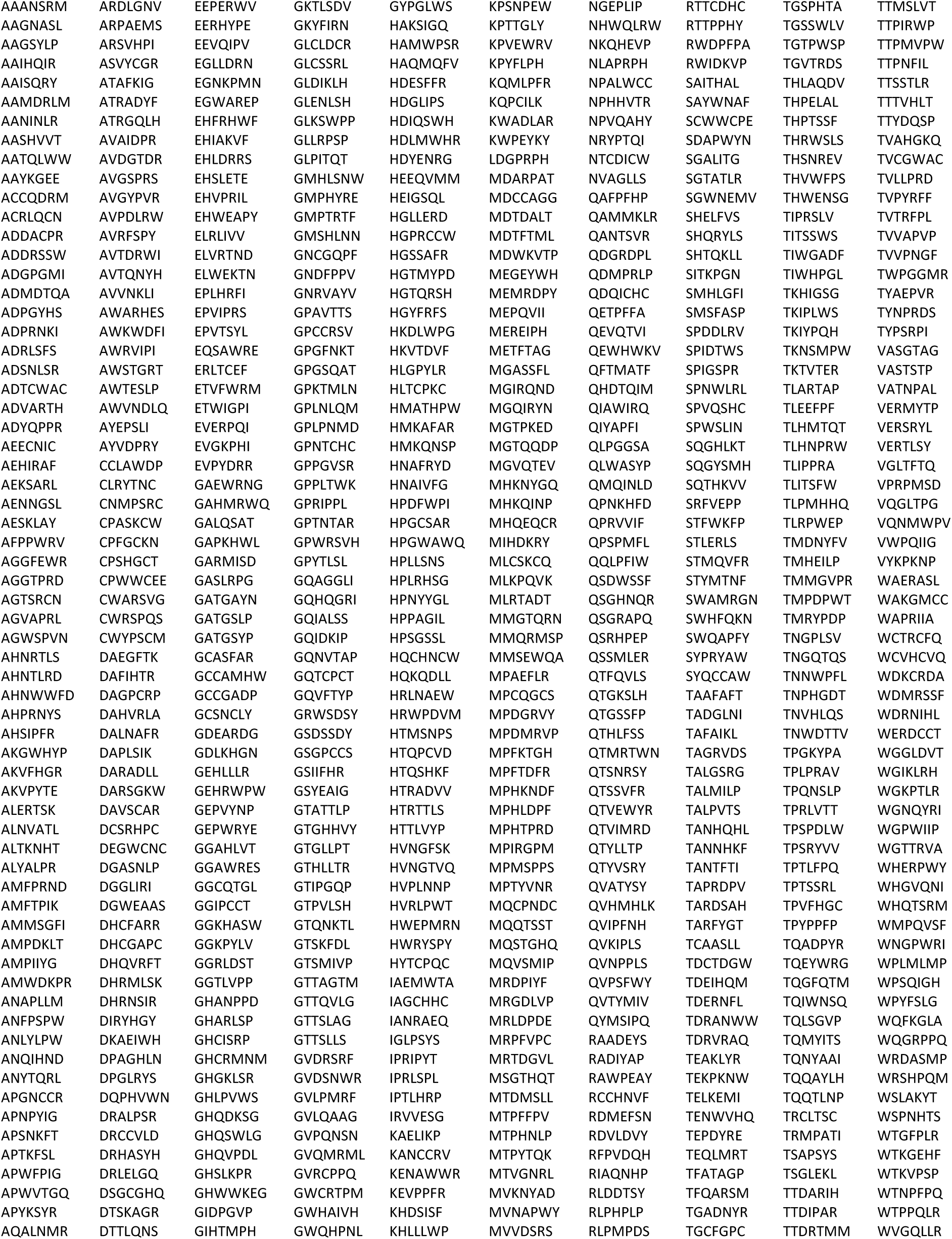
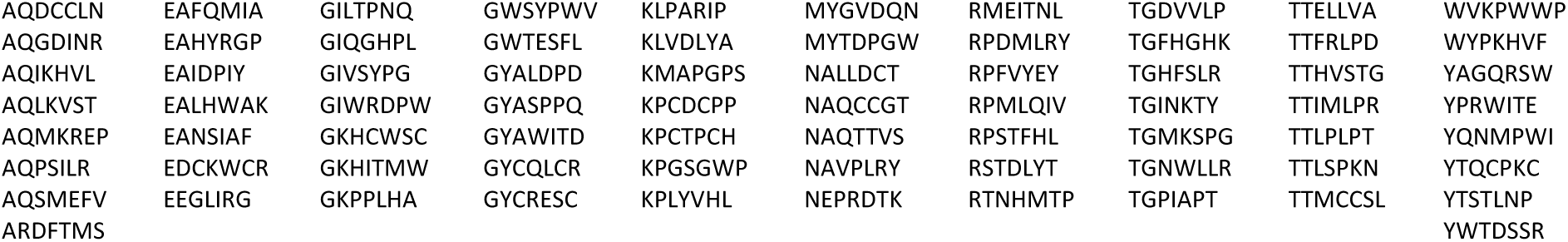
Sequences of the peptide mimotopes used in the study.

**Supplemental Table 2.**
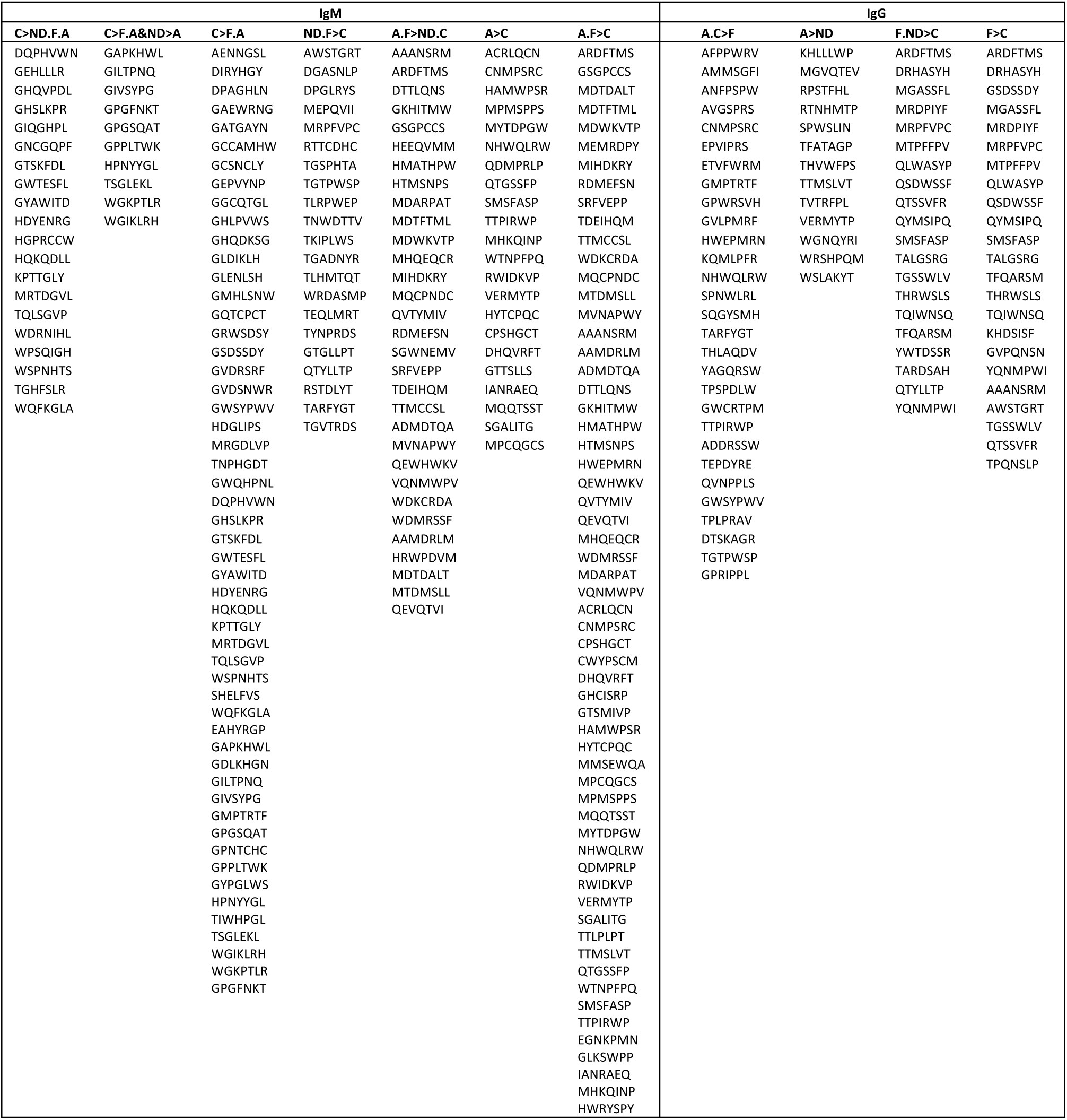
Mimotope clusters with significantly different expression in the different diagnoses after the final clustering.

**Supplementary Table 3.**
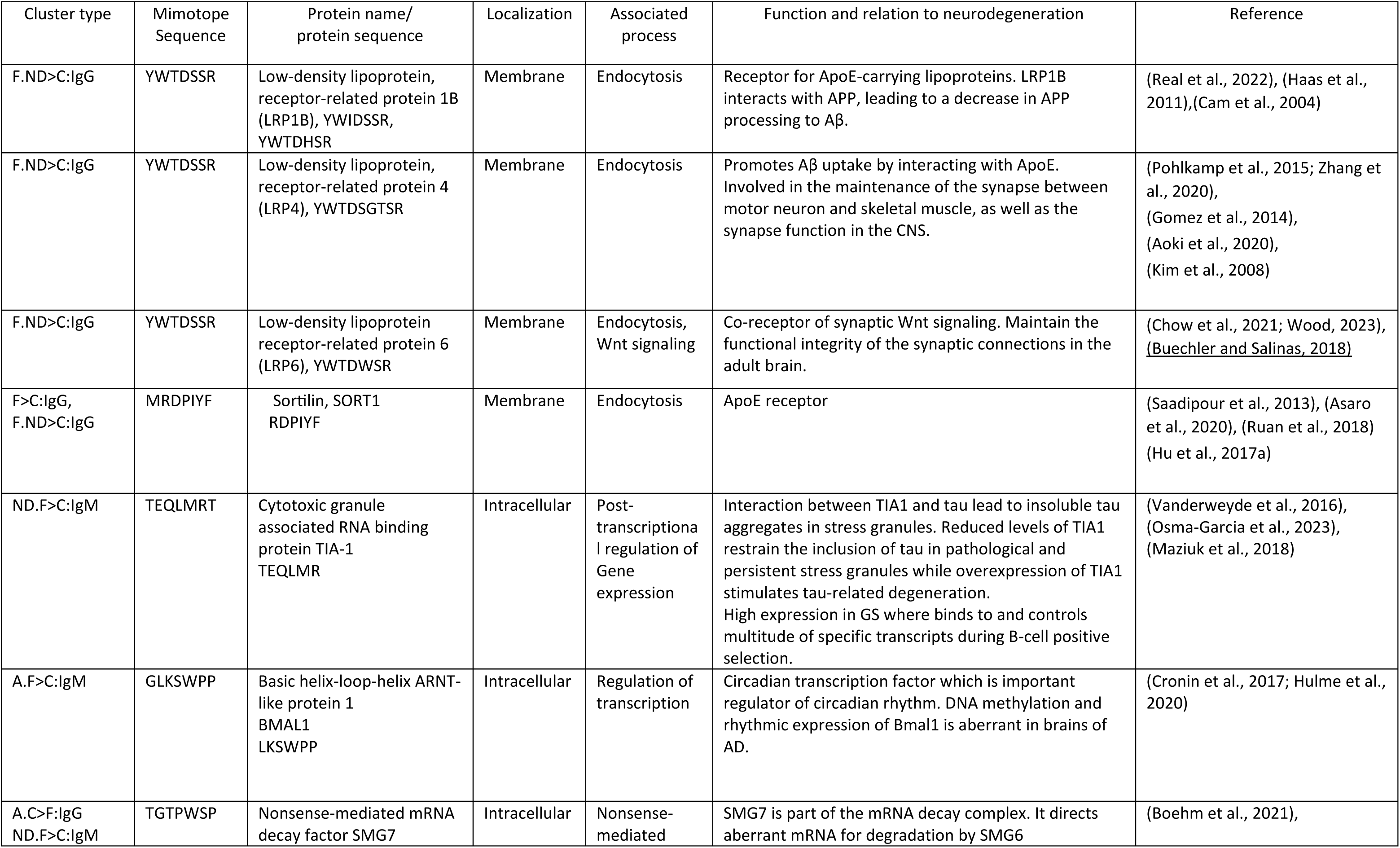
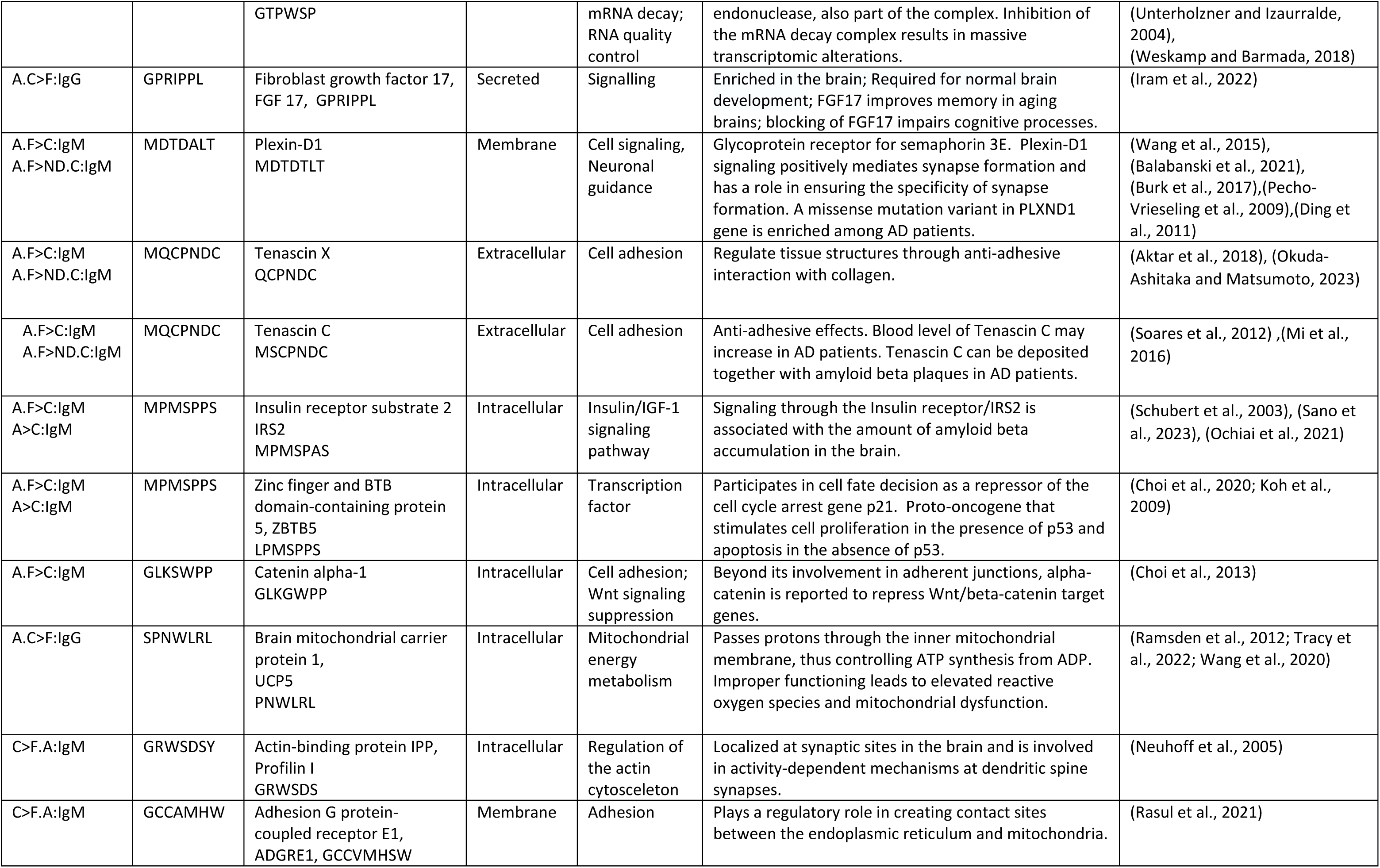
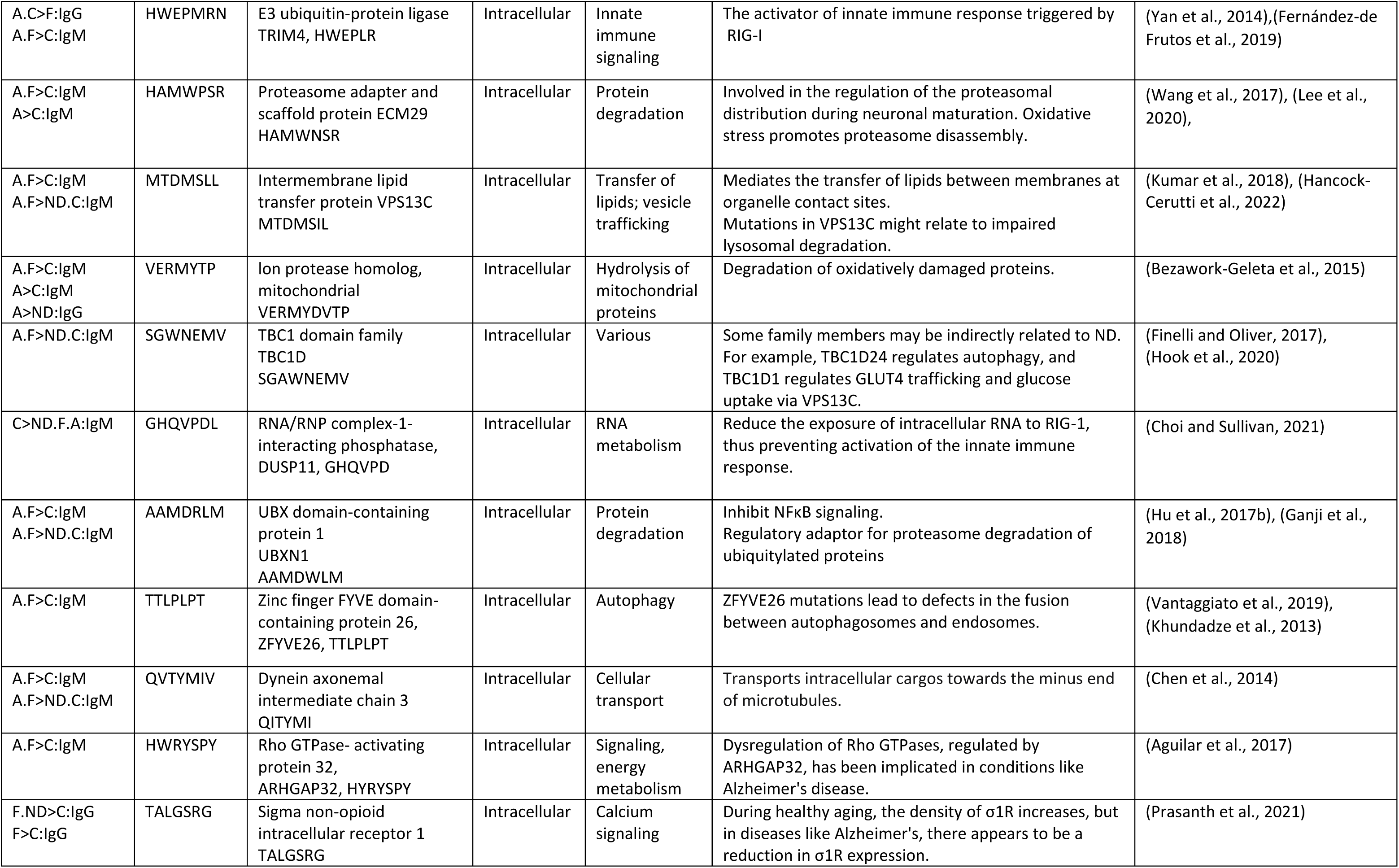
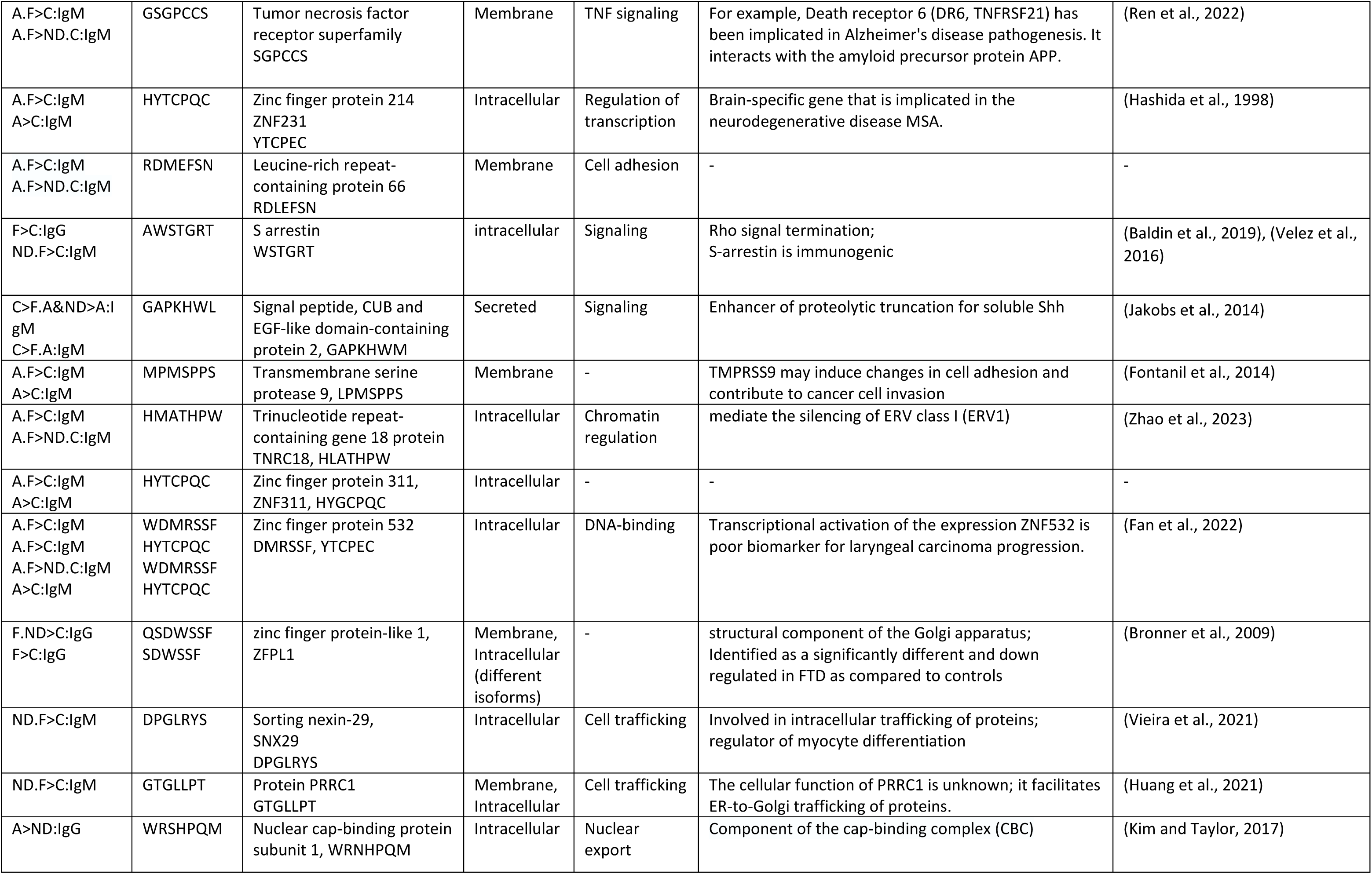
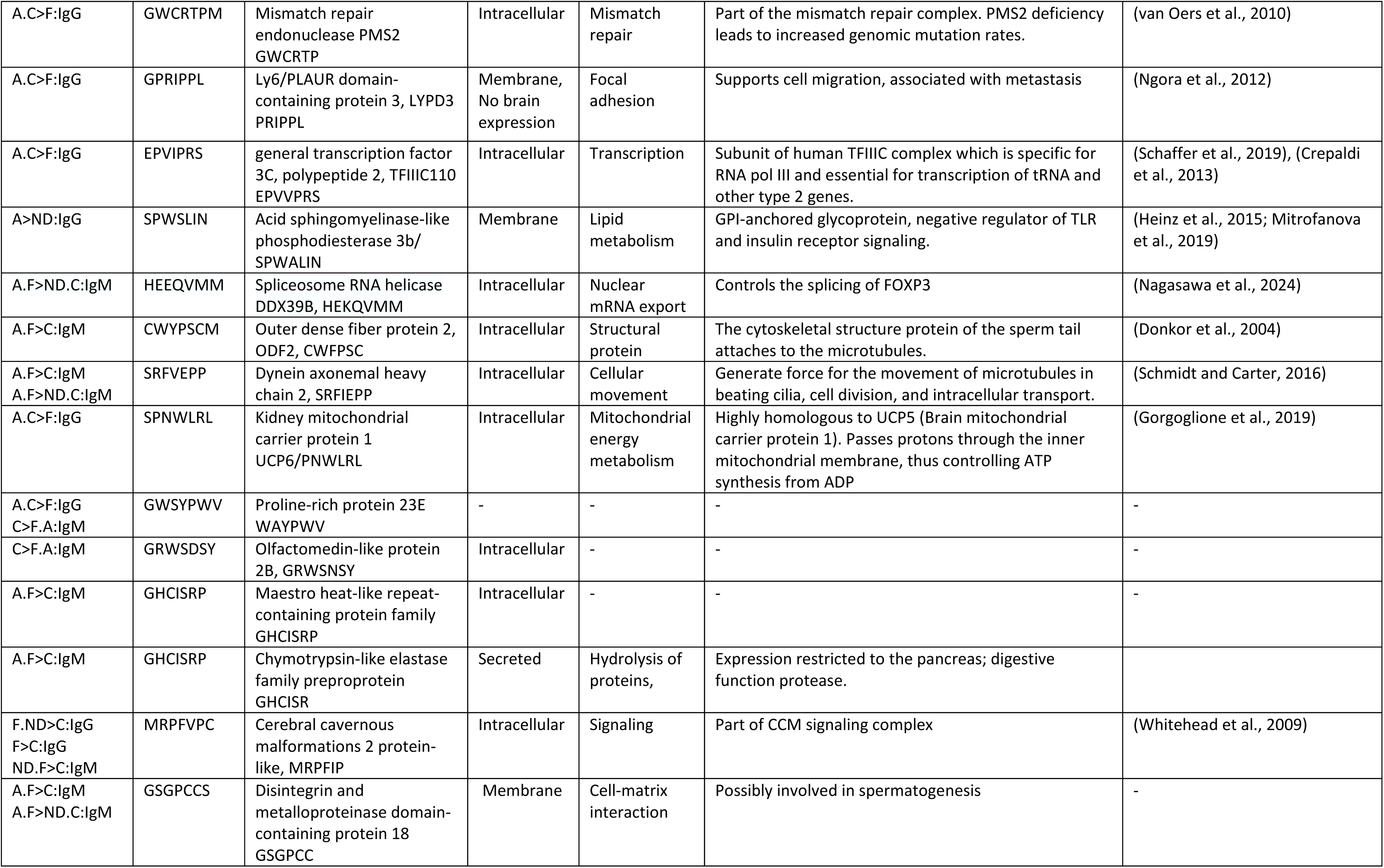
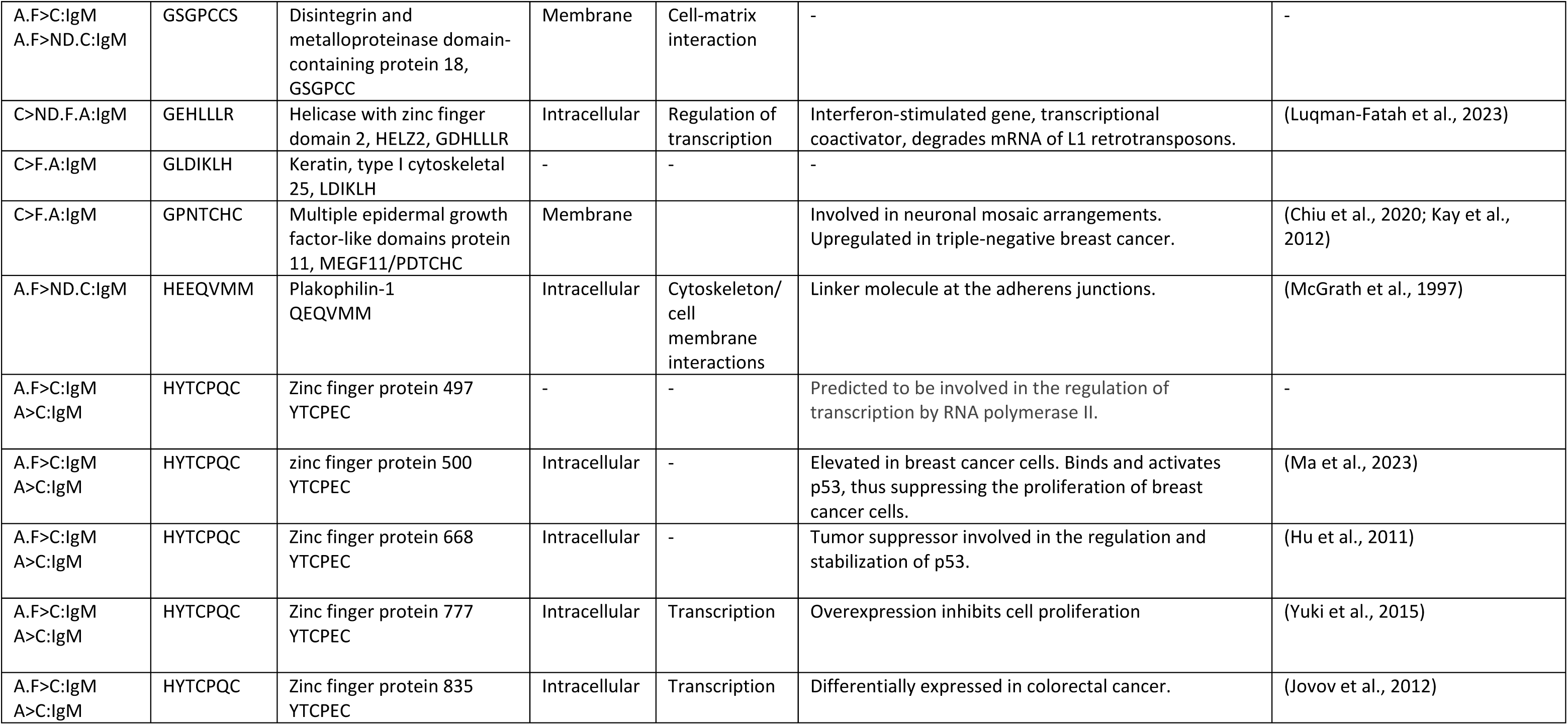
Proteins containing potential linear epitopes homologous to the mimotopes with differentially expressed antibody reactivities in AD and FTD.

**References to Suppl. Table 4.**
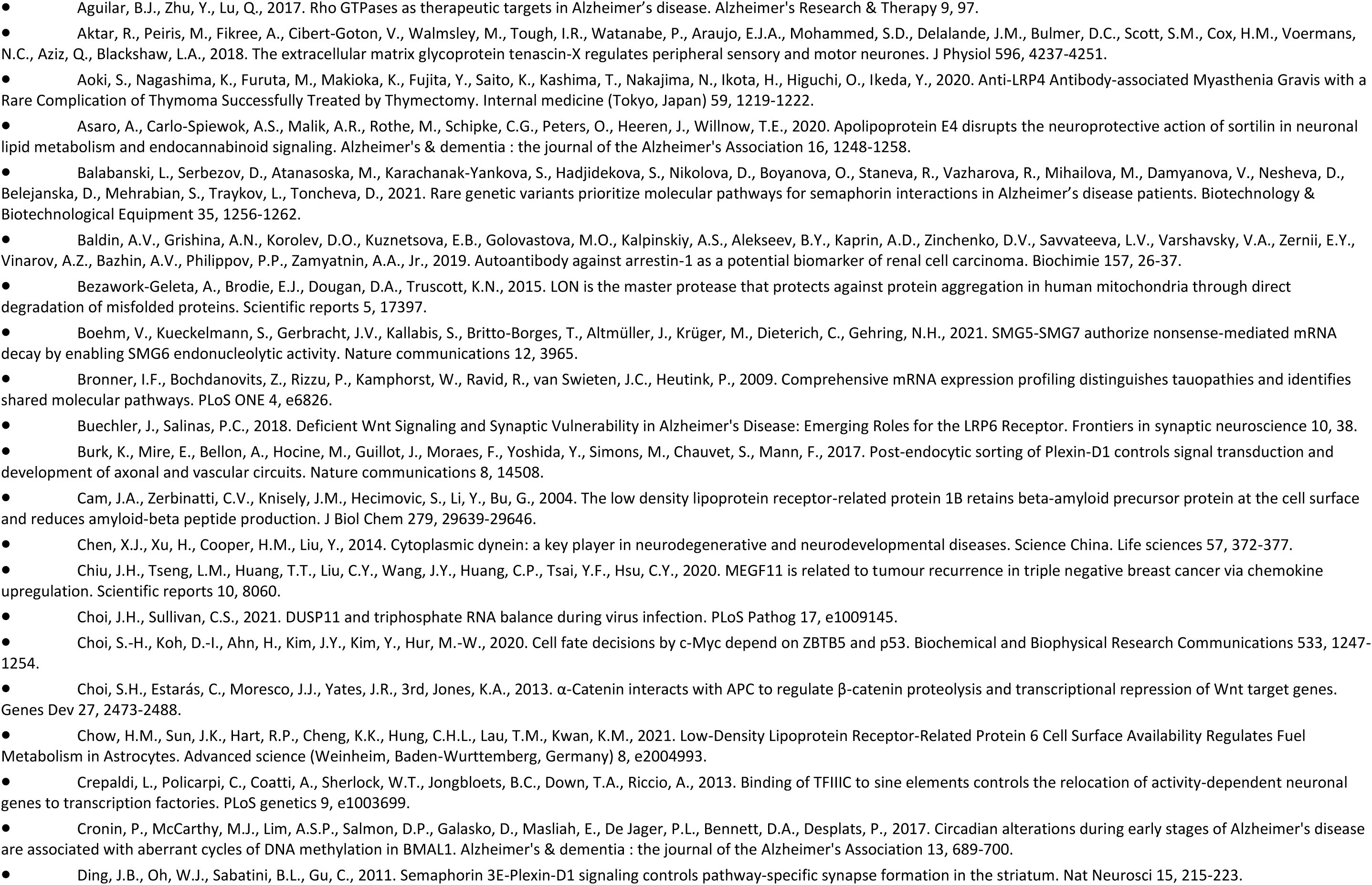
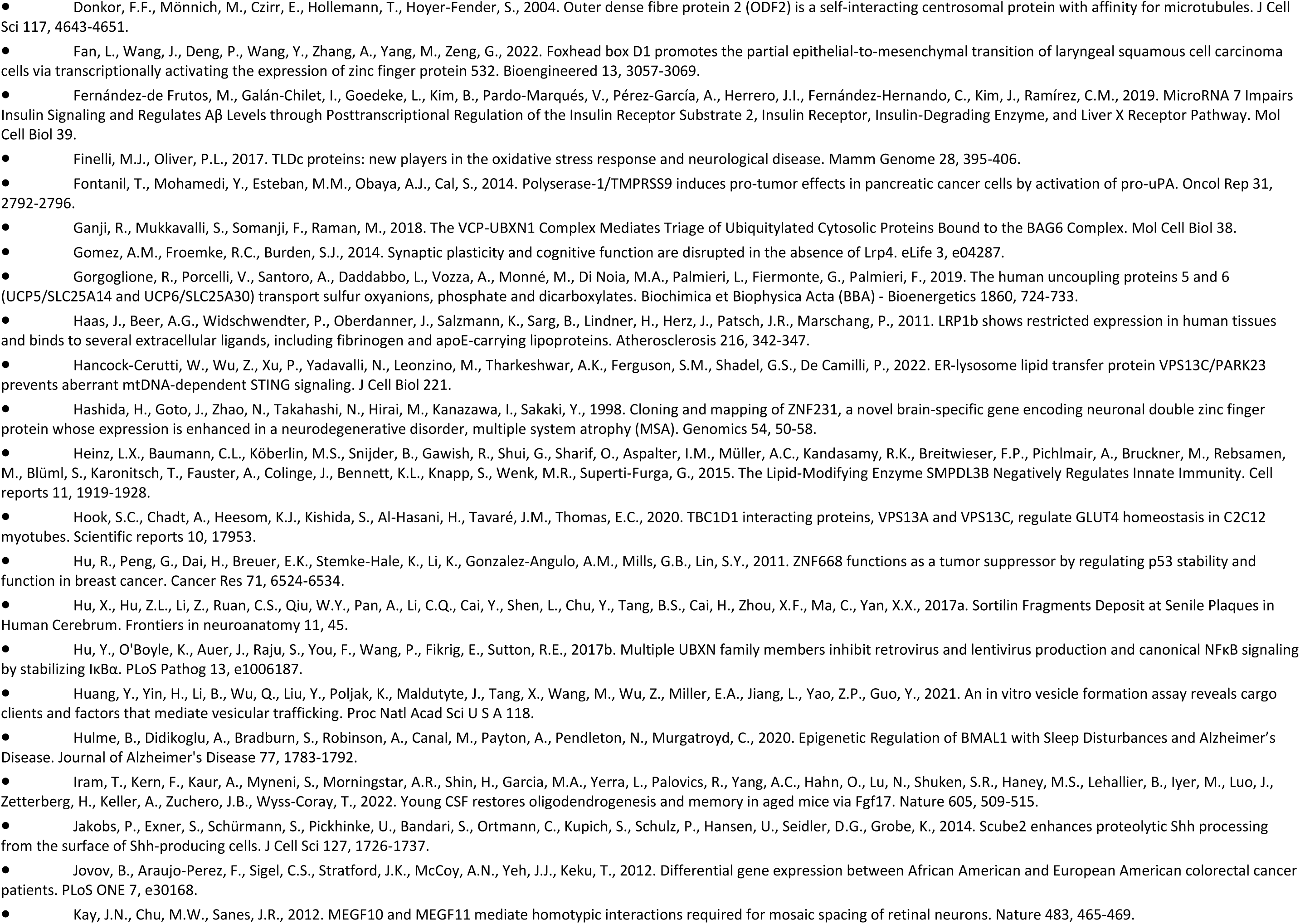
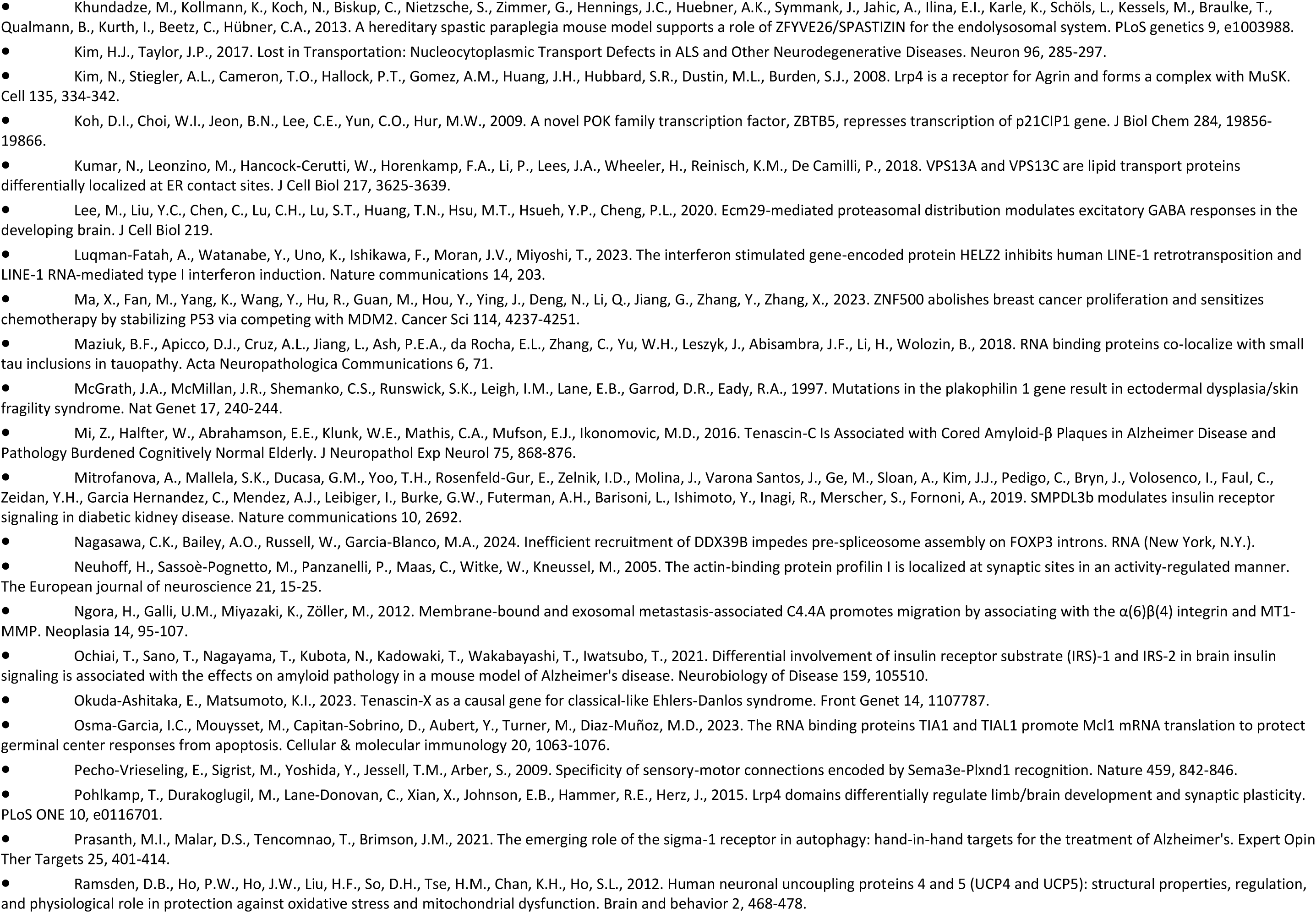
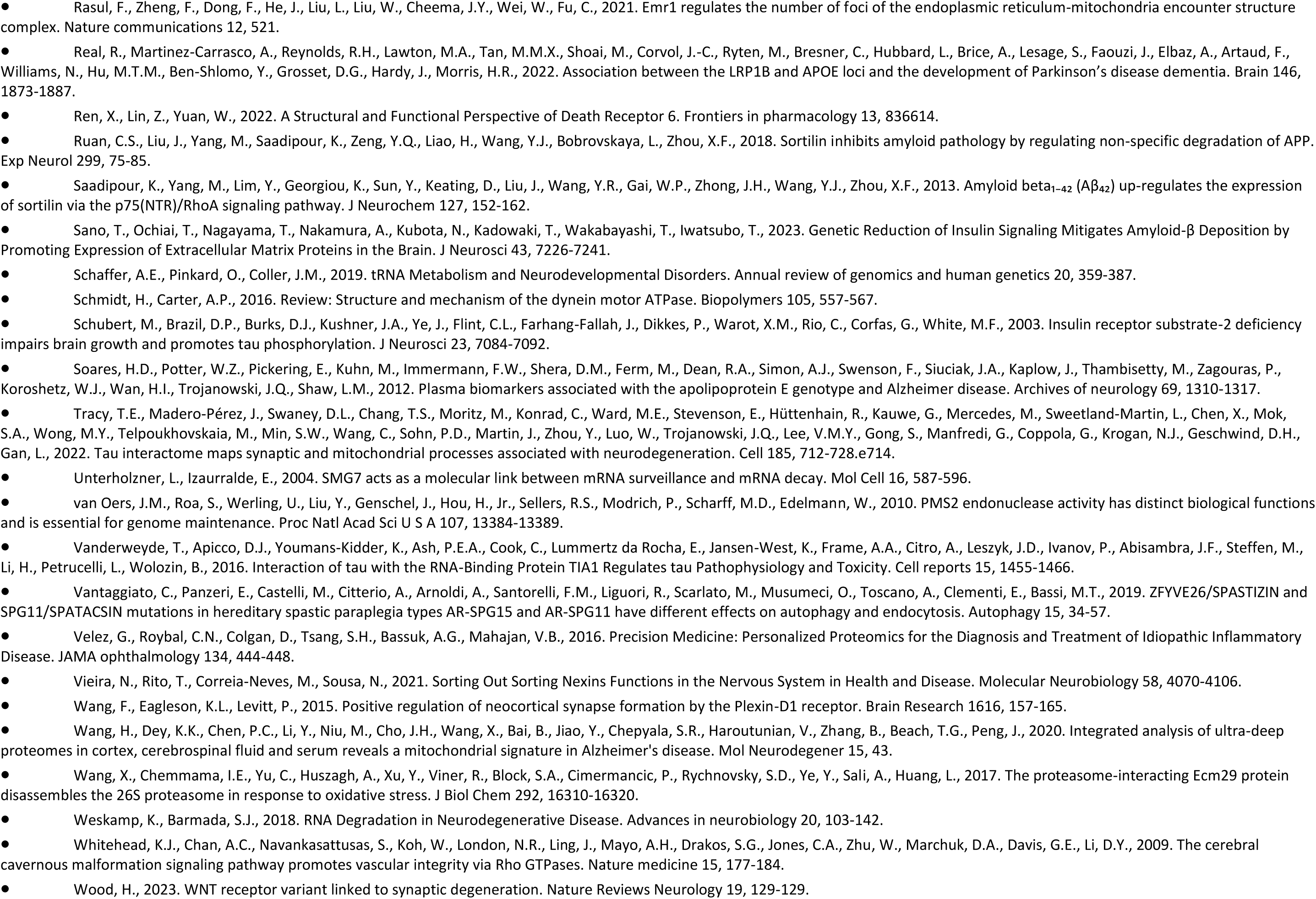
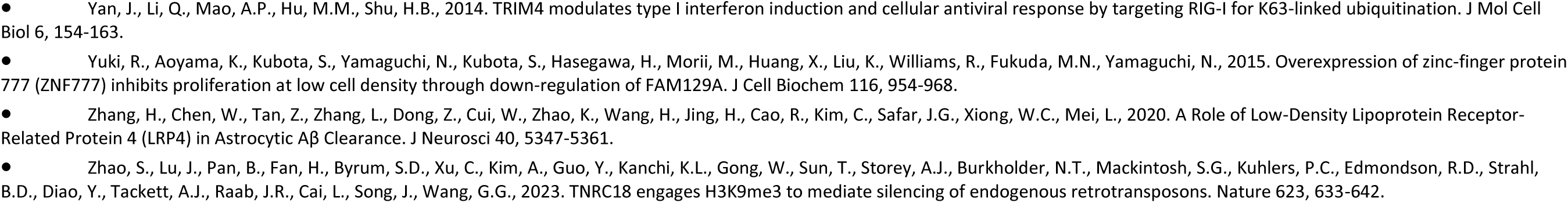

**Supplementary Figure 1.**
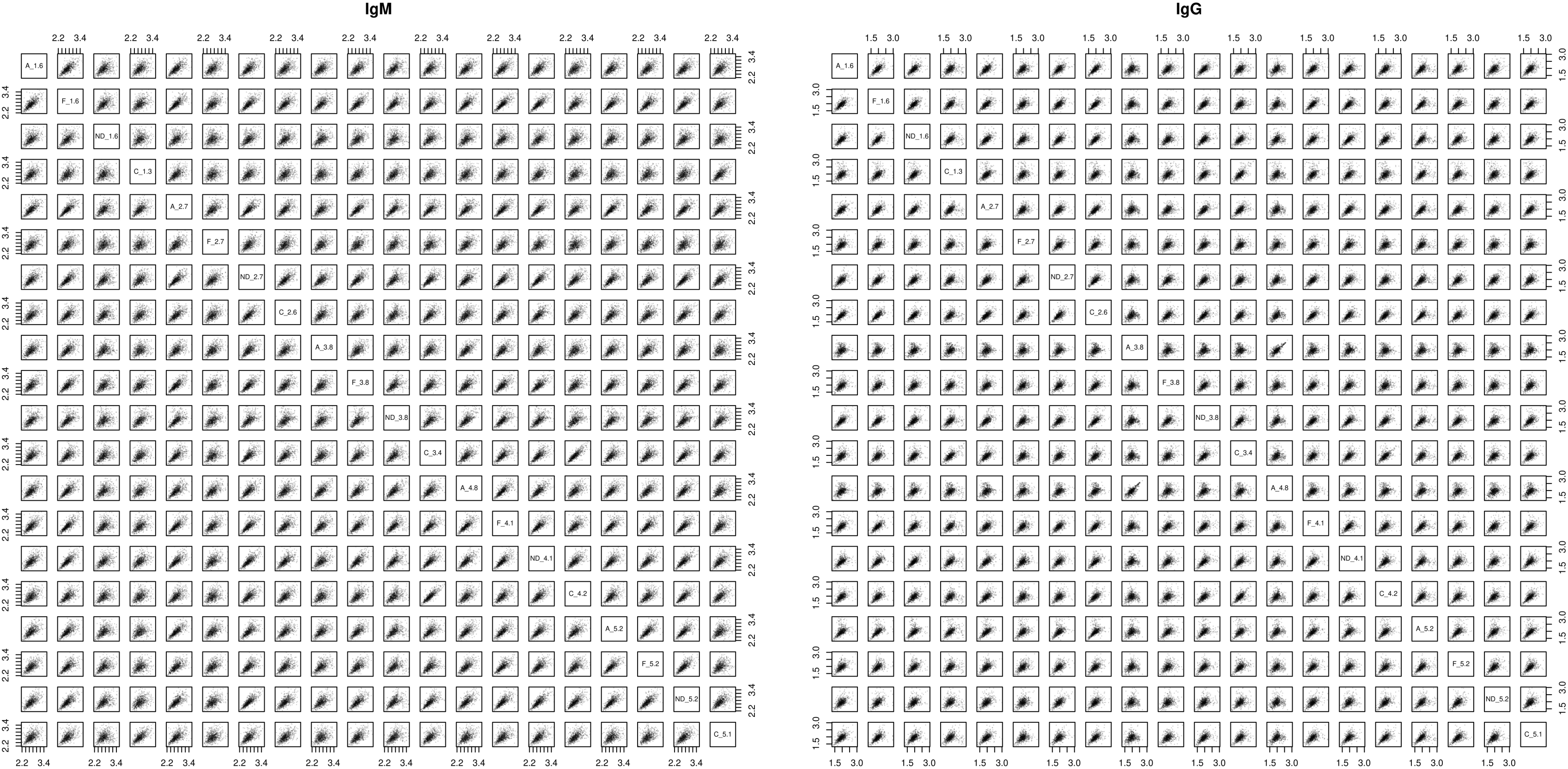
Pairwise scatter plots of the raw data after between array and compositional bias (dependence on charge, hydrophobicity, etc. general physicochemical properties of the residues) normalization.

**Supplementary Figure 2.**
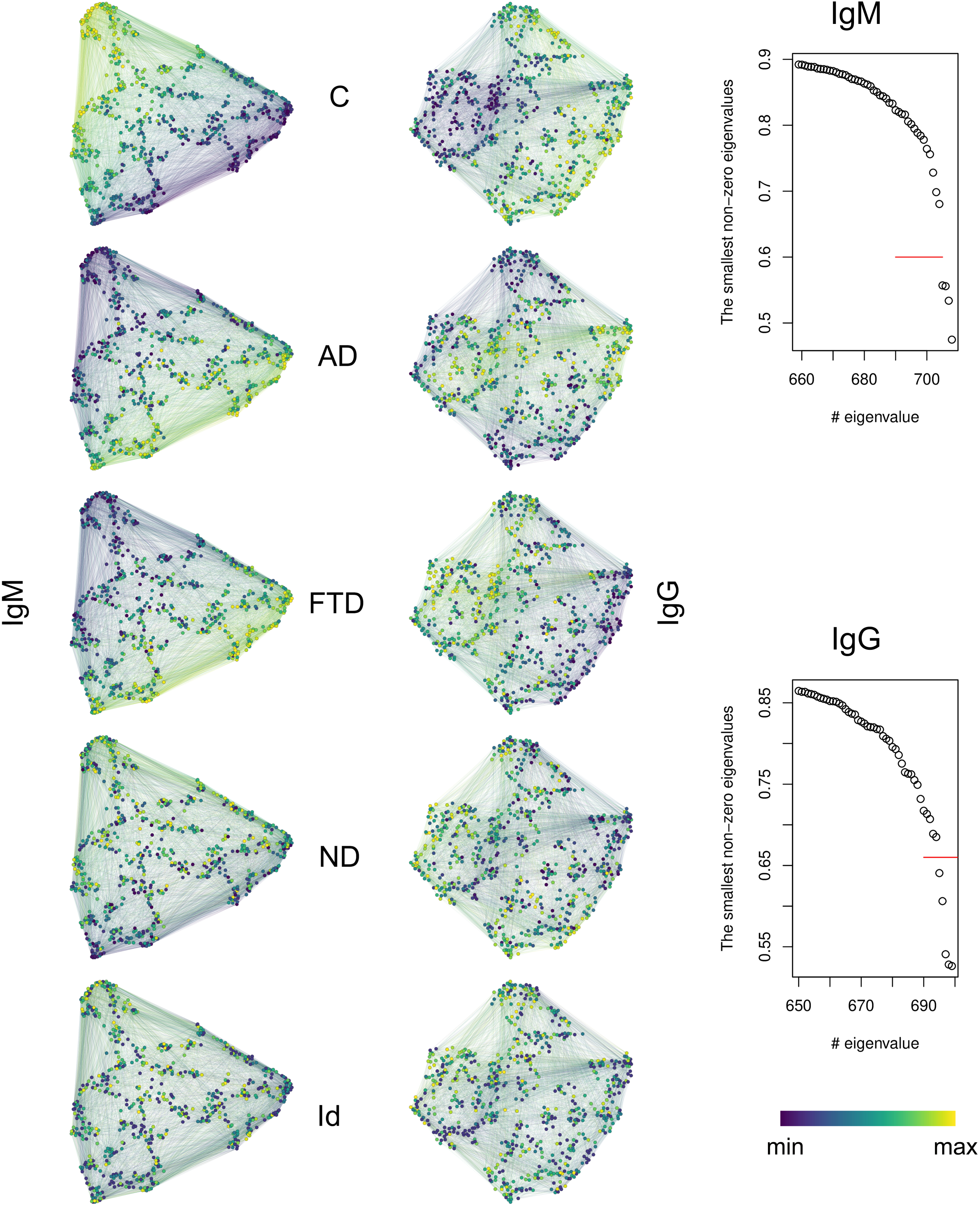
Cross-reactivity graph spectral embeddings for the IgM (left) and the IgG (center) graphs. The embedding uses the eigenvectors of the graph Laplacian that correspond to the smallest eigenvalues with a cut off based on irregular increase in the ordered eigenvalues (right) – 4 dimensions for IgM and five dimensions for IgG. The exact position of the vertices in the two-dimensional projections is derived by UMAP transformation from 4 (5) dimensions to two dimensions. The vertices are color-coded with high values corresponding to the high expression in the respective diagnosis. The values mapped to them in this way correspond to the levels of reactivity as sums of the ranks of the indicated samples by group (AD, FTD, ND, and control - C), or to the number of nearest neighbors among the idiotopes. The edges inherit their color from their incident vertices by averaging. A clear separation is observed between the reactivities higher in the C vs those higher in AD or FTD. ND related reactivities differ from all.

**Supplementary Figure 3.**
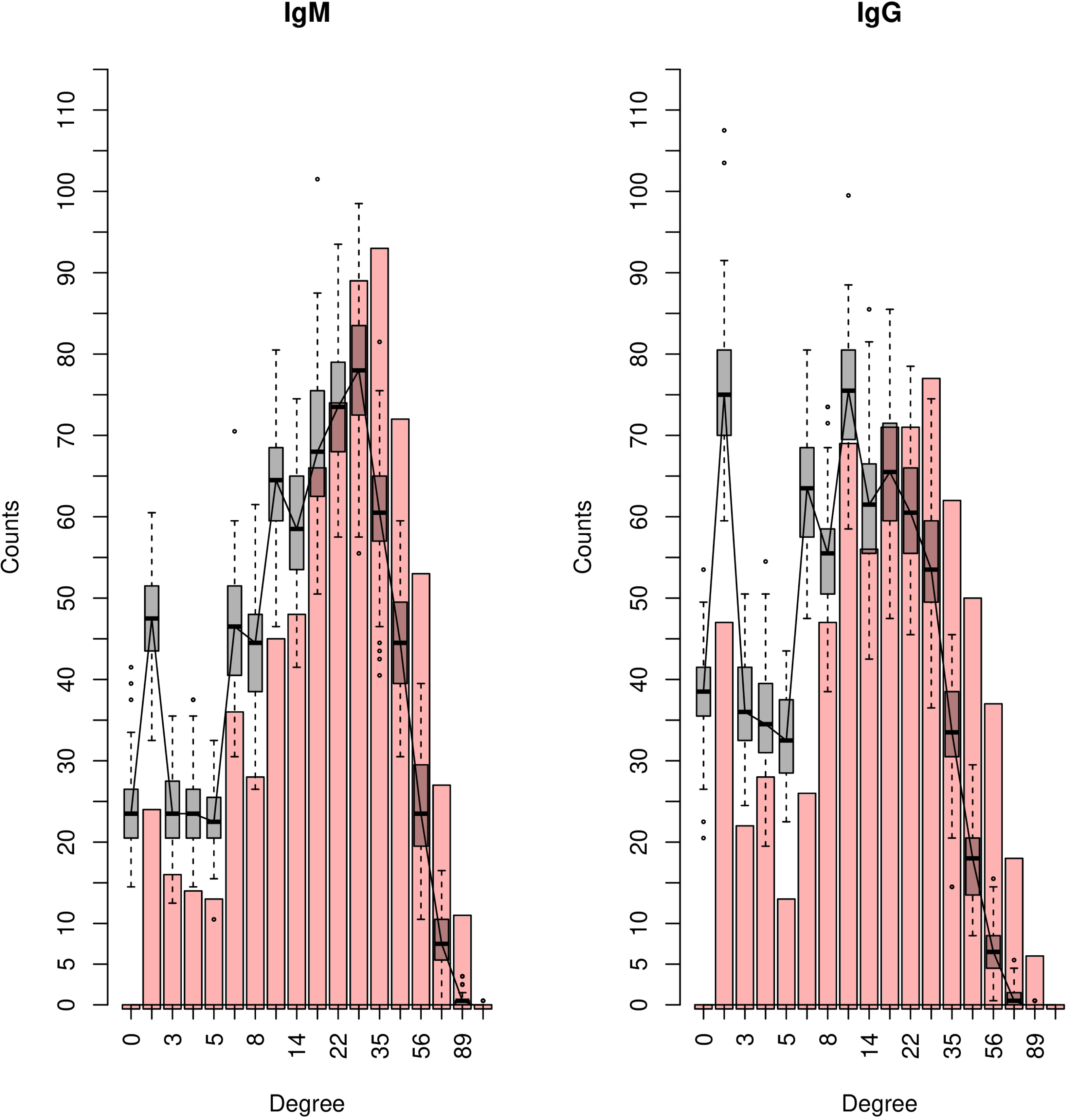
Degree distribution for the IgM/IgG graphs compared to the distributions of 100 graphs generated after scrambling the respective raw data matrices. The graphs have degree distribution drawn to the right, which corresponds to their higher the density as compared to randomly generated graphs.

**Supplementary Figure 4.**
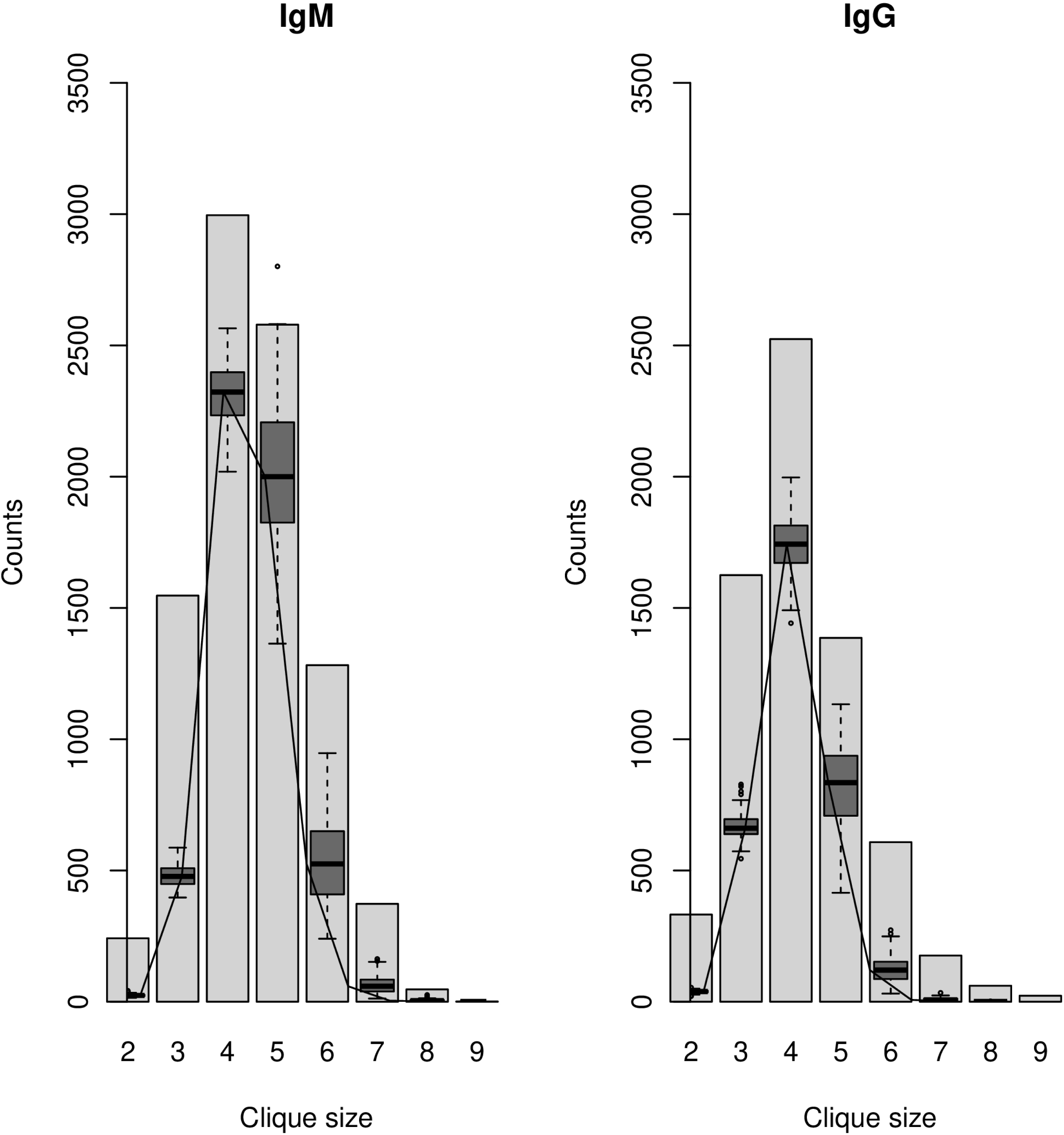
Maximal clique size distribution for the IgM and IgG graphs compared to the distributions of 100 graphs generated after scrambling the respective raw data matrices. The graphs have the same size but more cliques as compared to randomly generated graphs.

**Supplementary Figure 5.**
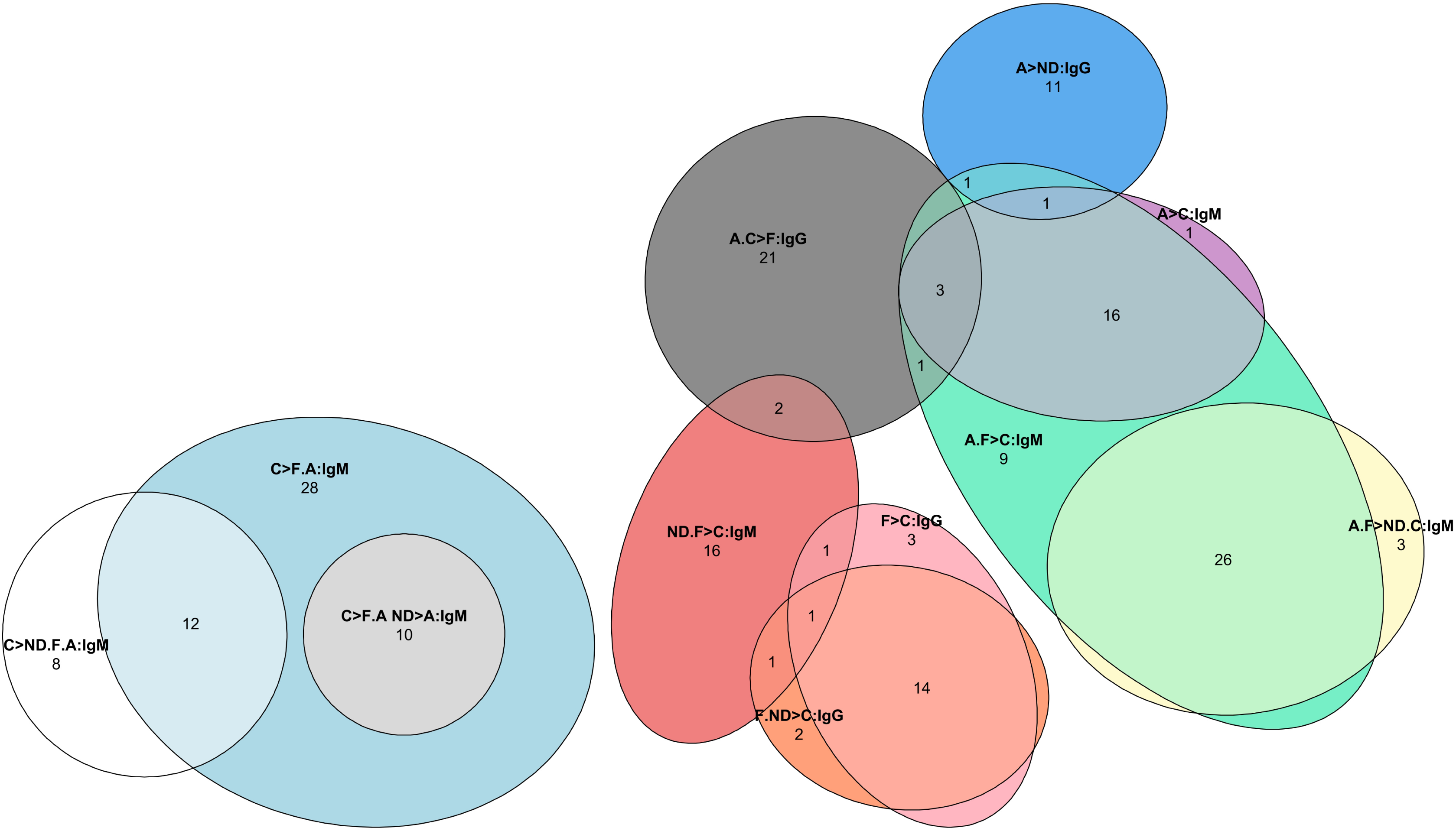
Venn diagram of the overlap between the ultimate cluster groups defined by the type of significant changes in the different diagnoses (annotation example: C>F.A ND>A:IgM – the reactivity in the controls is higher than that in FTD and AD plus NDD is only higher than AD, an IgM set clusters). The numbers indicate the number of mimotopes in this section. Some overlaps are between opposing characteristics, respectively from the IgM and the IgG sets (e.g. – some reactivities higher in FTD and NDD relative to the controls in IgG are found also in the group with the opposite ratios in IgM).

The supplementary files IgM_AD_FTD_NDD_C.mp4 and IgG_AD_FTD_NDD_C.mp4 visualize 3D embedding of the files in motion. The color code is mapping the rank sums by diagnosis AD, FTD and NDD scaled respectively as the red, green and blue components of the color while the luminosity codes for the reciprocal of the rank sum of the controls. Thus, the darker or black vertices correspond to reactivities found high in the controls but lost in the neurodegenerative diseases. In the IgM graph a considerable portion of the vertices with very light colors (reactivities not found in the controls) are colored red and green yielding a spectrum of predominantly yellow hues – the overlap of AD and FTD associated reactivities. In the IgG graph AD and FTD reactivities are separated in bright red and green clusters.

